# High-order theta harmonics account for the detection of slow gamma

**DOI:** 10.1101/428490

**Authors:** Y. Zhou, A. Sheremet, Y. Qin, J.P. Kennedy, N.M. DiCola, A. P. Maurer

## Abstract

Local field potential (LFP) oscillations are the superposition of excitatory/inhibitory postsynaptic potentials. In the hippocampus, the 20-55 Hz range (‘slow gamma’) is proposed to support cognition independent of other frequencies. However, this band overlaps with theta harmonics. We aimed to dissociate the generators of slow gamma versus theta harmonics with current source density and different LFP decompositions. Hippocampal theta harmonic and slow gamma generators were not dissociable. Moreover, comparison of wavelet, ensemble empirical-mode (EEMD), and Fourier decompositions produced distinct outcomes with wavelet and EEMD failing to resolve high-order theta harmonics well defined by Fourier analysis. The varying sizes of the time-frequency atoms used by wavelet distributed the higher-order harmonics over a broader range giving the impression of a low frequency burst (“slow gamma”). The absence of detectable slow gamma refutes a multiplexed model of cognition in favor of the energy cascade hypothesis in which dependency across oscillatory frequencies exists.

## Introduction

*In vivo* neurophysiology has been a major tool in attempting to understand how the brain organizes behavior, with local-field potential (LFP) oscillations taking center stage. The large rhythmic oscillation in the hippocampus, theta (4-12Hz), was among the first to be characterized in the freely behaving animal (Jung and Kornmüller, 1938; Green and Arduini, 1954; Green and Machne, 1955; Vanderwolf and Heron, 1964; Vanderwolf, 1969; for review, see Buzsáki, 2005). The smaller amplitude, faster gamma oscillation (Stumpf, 1965; Leung, 1992; Bragin et al., 1995; Chrobak and Buzsáki, 1998; Penttonen et al., 1998), initially defined as a single, broad frequency range (e.g., 40-100 Hz, Bragin et al., 1995), has recently been subdivided into two, sometimes three independent ranges referred to as slow-, medium- and fast gamma (Colgin et al., 2009; Belluscio et al., 2012; Carr et al., 2012; Colgin, 2012; Schomburg et al., 2014; Colgin, 2015; Fernández-Ruiz et al., 2017; Lopes-Dos-Santos et al., 2018; Michaels et al., 2018). While the exact definition of these sub-bands varies across publications, they have been generally identified as 20-55Hz (slow), 55-90Hz (medium), and fast (90-120Hz). The identification of more than one gamma band has led to the multiple-gammas hypothesis of cognition, which proposes that different gamma frequencies reflect different synaptic inputs to the hippocampus and possibly distinct cognitive processing states (for review, see Colgin, 2015). Notably, the data have been equivocal on what those states might be, with both slow and fast gamma being linked to memory retrieval processes (Shirvalkar et al., 2010; Kemere et al., 2013; Newman et al., 2013; Bieri et al., 2014; Igarashi et al., 2014; Schomburg et al., 2014; Takahashi et al., 2014; Trimper et al., 2014; Yamamoto et al., 2014). Moreover, the theta and gamma oscillations are not strictly memory-dependent. Rather, a substantial literature base demonstrates that these oscillations are velocity modulated (Chen et al., 2011; Ahmed and Mehta, 2012; Kemere et al., 2013; Zheng et al., 2015; Sheremet et al., 2018b).

Therefore, the exact relationship between the different gamma ranges and behavior is not straight forward. The problem is further complicated by the existence of theta harmonics (Harper, 1971; Coenen, 1975; Leung et al., 1982; Buzsaki et al., 1983; Leung and Buzsáki, 1983; Buzsáki et al., 1985; Ning and Bronzino, 1993; Czurkó et al., 1999; Terrazas et al., 2005), which are frequency integer, phase coupled oscillations that precipitate in spectral decomposition as a consequence of the fundamental 8Hz rhythm becoming both skewed and asymmetric. As the harmonics of theta have been observed to go as high as 48 Hz (Sheremet et al., 2016; Sheremet et al., 2018b), caution has been emphasized when examining the slow gamma band to avoid harmonic contamination (Schomburg et al., 2014; Scheffer-Teixeira and Tort, 2016). Specifically, theta harmonics (8, 16, 24, 32, 40, and 48 Hz) spill into the slow gamma range (25-50 Hz as defined by Colgin et al., 2009), potentially obfuscating spectral analyses.

In order to address the overlap between slow gamma and high-order theta harmonics in the power spectra, we sought to disambiguate the theta harmonics from slow gamma using current source density analysis across hippocampal lamina as well as spectral decomposition. Contemporary theories of slow gamma suggest that the oscillation is generated via CA3 to CA1 projections whereas the fast gamma is generated by entorhinal to CA1 projections (Colgin et al., 2009; Belluscio et al., 2012; Schomburg et al., 2014; Colgin, 2015; Hsiao et al., 2016; Zheng et al., 2016; Fernández-Ruiz et al., 2017). As these projections terminate in different layers of the hippocampus, str. radiatum and str. lacunosum-moleculare respectively, we hypothesized that a dissociation in the current source density pattern between these two oscillatory frequencies could be identified. Remarkably, in neither our own data nor data provided from another laboratory (Pastalkova et al., 2015; gift from the Buzsaki laboratory), were we able to find this dissociation. This concern was increased in our failure to detect slow gamma-theta phase coupling via bicoherence analysis which does not require any data preprocessing (Sheremet et al., 2016; Sheremet et al., 2018b). Furthermore, using the methods of Masimore and colleagues of determining the correlation coefficients between Fourier decompositions in order to identify fundamental frequencies and cross-frequency interactions (2004; 2005), coupling was only evident between theta harmonics and a single 50-120 Hz gamma band; there was a notable absence of slow gamma (Sheremet et al., 2018b). Finally, the absence of slow gamma was further supported by recapitulating the phase-amplitude analyses of Colgin et al. (2009) as a function of velocity (Sheremet et al., 2018b), which only revealed theta, theta harmonics and a broad 50-120Hz band coupling.

In light of these incongruencies, we revisited many of the major tenets that have led to the subdivision of the gamma band. Primary support for identifying the presence of an oscillation is an increase in power relative to background activity (i.e., the noise spectrum). Although initial approaches using Fourier based decompositions did not observe a deviation that is supportive of slow gamma (e.g., Buzsáki et al., 2003), wavelet decomposition (Colgin et al., 2009) and more recently, ensemble empirical mode decomposition (Lopes-Dos-Santos et al., 2018), indicate a power deviation in the slow gamma range. In investigating this, the methodological comparison suggest that slow gamma is an algorithmic artifact of convolving the harmonics of theta. As secondary analyses, such as spike modulation, coherence analyses, cross-regional interactions, and velocity modulation - *insufficient to determine the presence of an oscillation*- have been provided as support for slow gamma, we also revisit these results.

The major consequence of our research is that, as the subdivision of gamma has provided a major foundation to the ideas of neural multiplexing and spectral fingerprinting (Lisman, 2005; Siegel et al., 2012; Watrous and Ekstrom, 2014; Gu et al., 2015), the multiplexed model of information routing becomes significantly weaker. Rather, these data support an energy cascade model of hippocampal LFP in which low frequencies provide the power, and thus entrainment, of higher frequencies (Sheremet et al., 2018b; Sheremet et al., 2018a).

## Methods

### 0.1 Subjects and behavioral training

All behavioral procedures were performed in accordance with the National Institutes of Health guidelines for rodents and with protocols approved by the University of Florida Institutional Animal Care and Use Committee. A total of seven 4-10 months old Fisher344-Brown Norway Rats (Taconic) were used in the present study. This was a mixed sex cohort comprised of r530, r538, r539, r544, r695, r779, r782 in order to integrate sex a biological variable and begin to alleviate the disparity in research focused exclusively on males. Upon arrival, rats were allowed to acclimate to the colony room for one week. The rats were housed and maintained on a 12:12 light/dark cycle. All training sessions and electrophysiological recordings took place during the dark phase of the rats’ light/dark cycle. Training consisted of shaping the rats to traverse a circular track for food reward (45mg, unflavored dustless precision pellets; BioServ, New Jersey; Product #F0021). During this time, their body weight was slowly reduced to 85% to their arrival baseline. Once the rat reliably performed more than one lap per minute, they were implanted with a custom single shank silicon probe from NeuroNexus (Ann Arbor, MI). This probe was designed such that thirty-two recording sites, each with a recording area of 177 μm, were spaced 60 μm apart allowing incremental recording across the hippocampal lamina. In preparation for surgery, the probe was cleaned in a 4% dilution of Contrad detergent (Decon Contrad 70 Liquid Detergent, Fisher Scientific) and then rinsed in distilled water.

### 0.2 Surgical procedures

Surgery and all other animal care and procedures were conducted in accordance with the NIH Guide for the Care and Use of Laboratory Animals and approved by the Institutional Animal Care and Use Committee at the University of Florida. Rats were initially sedated in an induction chamber. Once anesthetized, the rat was transferred to a nose cone. The head was shaved with care taken to avoid the whiskers. The rat was then transferred to the stereotax, gently securing the ear bars and placing the front teeth over the incisor bar. The stereotaxic nose cone was secured, ensuring that the rat was appropriately inhaling the anesthesia. During surgical implantation, the rats were maintained under anesthesia with isoflurane administered at doses ranging from 0.5 to 2.5%. Next, ophthalmic ointment was applied and tanning shades, fabricated out of foil, were placed over but not touching the eyes to minimize direct light exposure. Multiple cycle of skin cleaning, using betadine followed by alcohol was applied prior to the first incision from approximately the forehead to just behind the ears. The remaining fascia was blunt dissected away and bone bleeding was mitigated through application of bone wax or cautery. Once the location of bregma was determined, the site of the craniotomy was located and a 3×3mm contour was drilled out, but not completed. This was followed by the placement of 7 anchor screws in the bone as well as a reference over the cerebellum and ground screw placed over the cortex. Once the screws were secured, a thin layer of adhesive cement (C&B-metabond, Parkell) followed by dental acrylic (Grip Cement Industrial Grade, 675571 (powder) 675572 (solvent); Dentsply Caulk, Milford, DE) were applied taking care to not obscure the craniotomy location. Finally, the craniotomy location was completed, irrigating and managing bleeding as necessary once the bone fragment was removed. Next a portion of the dura was removed, taking care to avoid damaging the vessels and the surface of the neocortex. Small bleeding was managed with saline irrigation and gel foam (sterile absorbable gelatin sponges manufactured by Pharmacia & Upjohn Co, Kalamazoo, MI; a division of Pfizer, NY, NY). The probe implant coordinates targeted the dorsal hippocampus (AP: −3.2 mm, ML: 1.5 relative to bregma, DV: −3.7 to brain surface). Some of these rats also received an identical 32- site probe targeting the medial entorhinal cortex (0.5mm anterior to the transverse sinus, ML: - 4.6mm, DV: −5.78mm from brain surface, angled 15° posterior). Once the probe was in place, the craniotomy was covered with silastic (Kwik-Sil, World Precision Instruments, Sarasota, FL) and then secured to the anchor screws with dental acrylic. Four copper mesh flaps were placed around the probe providing protection as well as acting as a potential Faraday cage. The wires from the reference and ground screws were soldered to the appropriate pins of the connector. Adjacent regions of the copper-mesh flaps were soldered together to ensure their electrical continuity and the ground wire soldered to the copper mesh taking care to isolate the reference from contact with the ground. Once the probe was secured, the rat received 10cc of sterile saline as well as metacam (1.0 mg/kg) subcutaneously (the non-steroidal anti-inflammatory is also known as meloxicam; Boehringer Ingelheim Vetmedica, Inc., St. Joseph, MO). The rat was placed in a cage and monitored constantly until fully recovered. Over the next 7 days, the rat was monitored to ensure recovery and no behavioral anomalies. Metacam was administered the day following surgery as well. Antibiotics (Sulfamethoxazole/Trimethoprim Oral Suspension at 200mg/40mg per 5 mls; Aurobindo Pharma USA, Inc., Dayton, NJ) were administered in the rat mash for an additional 5 days.

### 0.3 Neurophysiology

Following recovery from surgery, rats were retrained to run unidirectionally on a circle track (outer diameter: 115 cm, inner diameter: 88 cm), receiving food reward at a single location. For rats 530, 544 and 695, data is only analyzed from the circle track conditions. In order to deal with low velocities from the circle track datasets, additional datasets for rats 538 and 539 from running on figure-8 track (112 cm wide × 91 cm length) were used. In this task, rats were rewarded on successful spatial alternations. Only datasets in which the rats performed more than 85% of trials correctly were used. The local-field potential was recorded on a Tucker-Davis Neurophysiology System (Alachua, FL) at 24 kHz (PZ2 and RZ2, Tucker-Davis Technologies). The animal’s position was recorded at 30 frames/s (Tucker-Davis). Spatial resolution was less than 0.5 cm/pixel. Speed was calculated as the derivative of the smoothed position. The local-field potential data was analyzed in Matlab^®^ (MathWorks, Natick, MA, USA) using custom written code as well as code imported from the HOSAtoolbox. Raw LFP records sampled at 24 kHz (Tucker-Davis system) were low-pass filtered down to 2 kHz and divided into fragments of 2048 time samples (approximately 1 second). To eliminate the effects of anatomic variations between individuals and because the position and orientation of the hippocampal LFP probes cannot be controlled precisely, the position of the hippocampal layers with respect to the recording channels was determined by estimating the distribution of current-source density (Rappelsberger et al., 1981; Mitzdorf, 1985; Buzsaki et al., 1986; Bragin et al., 1995). Current-source density exhibits a unique distribution that is largely independent of anatomic details and that may be used to accurately identify hippocampal layers.

#### Spike autocorrelation and cross-correlation spectrogram analyses

Often the presence or absence of an oscillation is buttressed by whether or not single units are modulated at a specific frequency, accomplished for examining if the spike times of the individual cells prefer a specific phase of an oscillation. However, nonlinear issues can become a factor in these analyses (Belluscio et al., 2012), necessitating a different approach. Therefore, to determine the frequency in which neuronal spiking occurred, we implemented spectral analyses on spike trains (Leung and Buzsáki, 1983; Sheremet et al., 2016). Single unit data was generously provided by the Buzsaki laboratory and curated by the Collaborative Research in Computational Neuroscience (Mizuseki et al., 2014; Pastalkova et al., 2015; datasets: i01_maze06.005, i01_maze08.001, and i01_maze08.004). For these dataset, the rat performed a delayed alternation task on a figure-8 maze, running in a wheel during the delay. Only bilaterally recorded CA1 neurons were used for the analysis. Action potentials for pyramidal cells and interneurons were initially sorted in to velocity bins, analyzing the 5-15 or 35+ cm/sec conditions separately. These spike times were converted into a binary time series with a sampling frequency of the 1250 Hz. As this binsize is slightly smaller than the traditional window of 1 msec used for waveforms, a box car convolution was performed such that a single spike registers as 1 across 3 adjacent bins. Each spike train was then passed through Matlab^®^’s xcorr.m function which finds the correlation at specified lag times. Lags we only calculated up to 3 seconds. Each output was then passed through the pmtmPH (https://www.mathworks.com/matlabcentral/fileexchange/2927-pmtmph-m) function to calculate the power spectral density (PSD) based on multitaper analysis. Each unit’s power was normalized by the total power and then sorted by peak theta frequencies to align the harmonics between units. The average and standard error power spectra is presented.

### Fourier and Wavelet Transforms

#### I. Transforms

The Fourier and wavelet transforms can be introduced formally as decomposition on an orthogonal basis. Let 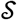 be some class of real functions and let *ψ_f_* (*f* ∈ *R*) be a basis in 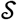; then any 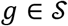 can be written uniquely as

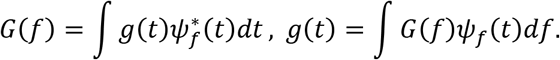

The “coefficients”*G*(*f*) of the decomposition (also referred to as the transform of *ɡ*) are obtained by taking the inner product of the function *ɡ* with the basis elements (i.e., projecting *ɡ*(*t*) onto the basis), The pair of equations are best known as direct and inverse transforms, sometimes also called analysis and synthesis.

The Fourier transform pair is obtained letting *ψ_f_*(*t*) = *e*^2*πift*^

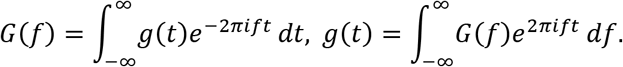

Unless the set of functions is carefully defined, these equations are only formal, and in most cases, they do not have any elementary mathematical meaning; when they do, their usefulness is limited. For example, if *ɡ*(*t*) = 1, the Riemann integral in the equation does not exist; however, a rather restrictive elementary theory can be built for *T*-periodic functions.

The wavelet transform and its inverse are given by

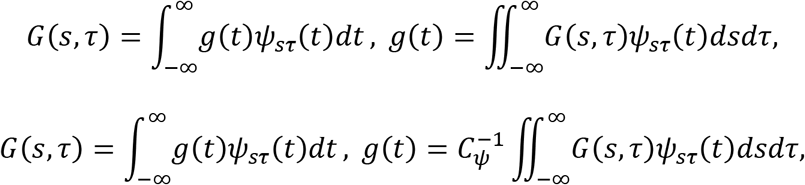

where the constant *C_ψ_* is the normalization constant

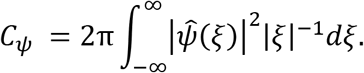

The functions 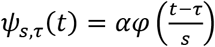 are “copies” of the “mother wavelet” *φ*(*t*), shifted in time by *τ* and scaled by *s*, with *α* a normalization constant. And 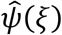 is the frequency space representation of the mother wavelet. The full set of wavelets {*ψ*_*s,τ*_(*t*)}_*s,τ*∈*R*_ is in general complete but not independent (i.e., larger than a basis).

#### II. Discrete Transforms

Evolving the generic Fourier and Wavelet equations into useful analysis tools has complexities that required the development of full theories (e.g., the theory of distributions, or generalized functions, e.g., Schwartz, 1950; Lighthill, 1958). For example, in Fourier equations *ψ_f_*(*t*) = *e*^2*πift*^ are orthogonal in the sense that 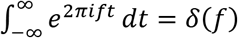, where *δ* is the Dirac delta function (Vladimirov, 2002; Strichartz, 2003). A complete discussion of these is far beyond the scope of this study, and is also unnecessary, because time series measured in practical applications are always of finite length *T*, sampled at time intervals Δ *t* (*T* = *n*Δ*t*), i.e., are finite sequences of real numbers *ɡ*(*t*) = {*ɡ*_1_, *ɡ*_2_,…, *ɡ*_*N*_, with *ɡ*_*j*_ = *ɡ*(*t_j_*). Such sequences naturally form *N*-dimensional spaces, in which integral Transforms, such as the Fourier or Wavelet equations, are represented by finite-dimensional linear operators, i.e., *N* × *N* matrices. These discretized versions of the Fourier and wavelet representations are called the *discrete* transforms (Briggs, 1995; Mallat, 1999; Strang, 2006).

The discrete Fourier transform pair is (in Matlab^®^ convention)

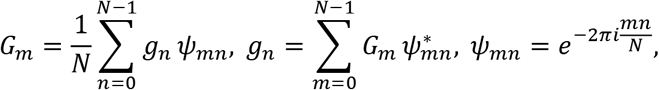

where 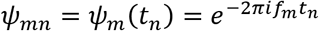 are the basis vectors. This pair of equations are sometimes called the analysis and synthesis of the signal *ɡ*(*t*).

Here, *f*_*m*_ = *m*Δ*f* and *t*_*n*_ = *n*Δ*t* represent the discretized frequency and time grids, with 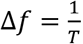, and 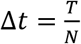. Basis functions are orthogonal in the sense that, for any integer *m*, with *m* ≠ 0 and *m* ≠ *N*, 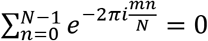.

A discrete version of wavelet transform is

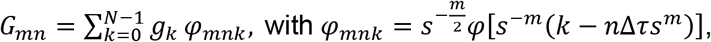

where Δ*τ* is a time-shift increment. The wavelets *φ_mnk_* form orthogonal only for compact-support wavelet shapes *φ* (e.g., Haar, and Daubechies wavelets; Daubechies, 1988, 1992; Mallat, 1999).

The Parseval relation insures that the discrete Fourier equations conserve the variance of the time sequence and its transform (e.g., Briggs, 1995), i.e., *σ*^*ɡ*^ = *σ*^*G*^, where *σ*^*ɡ*^ = ∑_*n*_|*ɡ*_*n*_|^2^ is the variance of *ɡ*. The discrete wavelet transform does conserve variance (*σ*^*ɡ*^ ≠ *σ*^*G*^) and in general, the variance ratio depends on *ɡ*, which means that a universal correction factor does not exist.

### Windowed Fourier Transform and the Correlation Coefficients of the Spectrogram

To investigate the evolution of spectra over time, windowed Fourier transform (WFT) is introduced where the original signal is multiplied by a window function which is nonzero for only a short period of time

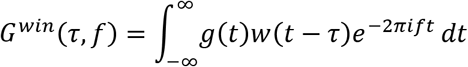

where *w* is the window function (Priestley, 1981; Roads, 2004). The window function slides along time axis, which gives rise to a time-frequency representation of the original time series. This time-frequency representation can be plotted as the spectrogram.

Typically, window functions are smooth, “bell shaped”, time-localized curves, the Fourier transform of the window function is

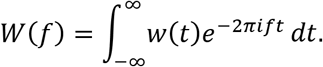

In frequency space, the window function *w* is usually composed of a “bell shaped” main lobe and symmetric side lobes. The window bandwidth of a window function is defined as

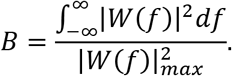

The spectrum estimation depends critically on the window bandwidth as it will smooth the spectrum and influence the frequency resolution (**Fig. 1**). The time-frequency resolution of transforms will be further discussed in the following section.

**Figure 1:**
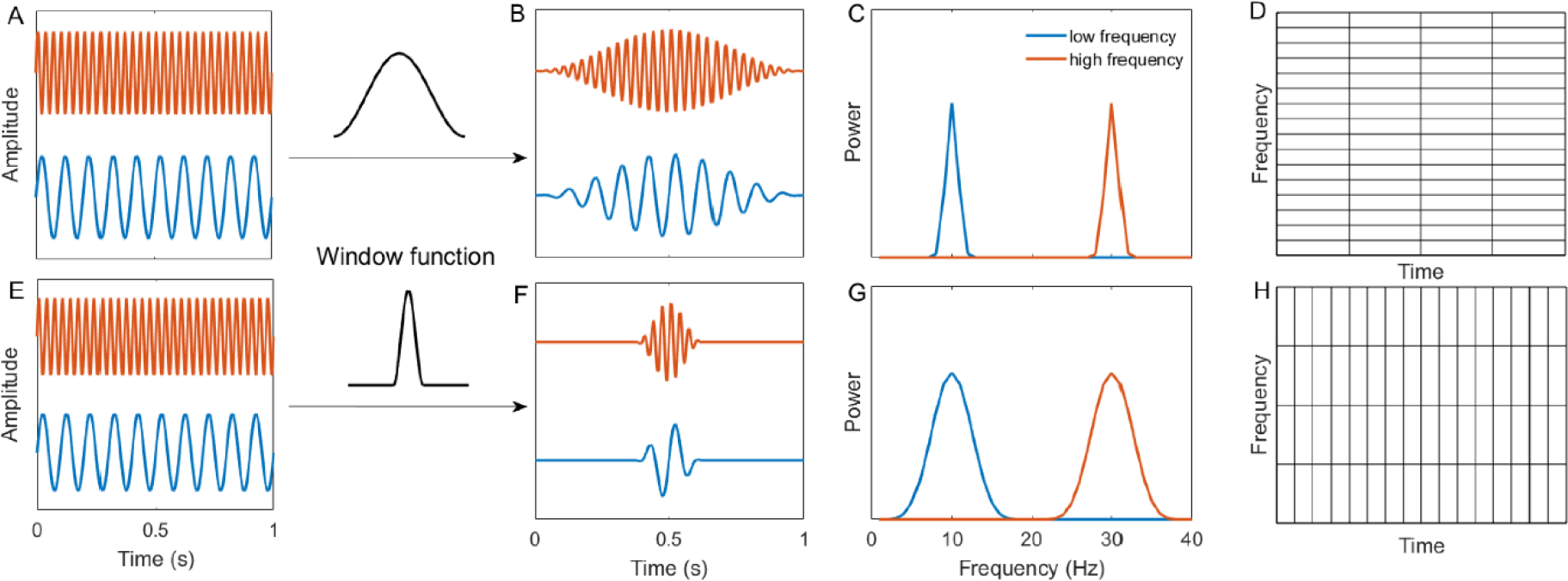
Windowed Fourier transform atoms. For windowed Fourier transform, window function is applied to all the frequency components. Adopting a Hanning window with wide time support to fast and slow oscillations (A) gives rise to fast and slow oscillations which decay slowly in time domain (B). In frequency domain, the power (amplitude square) of these slowly decay oscillations have relatively narrow frequency band and the bandwidth is the same for fast and slow oscillations (C). This results in transform with coarse time resolution but fine frequency resolution (D). As a result, windowed Fourier transform with wide time supported window functions has fine frequency resolution but coarse time resolution. On the contrary, applying Hanning window with narrow time support leads to fast decay oscillations in time domain (F). However, these fast decay oscillations have wide frequency band (G) and result in transform with fine time resolution but coarse frequency resolution (H).

In order to approach LFP interactions across regions, we implemented the method of calculating the correlation coefficients of the spectrogram as outlined by Masimore and colleagues (2004; 2005). The power cross-correlograms were obtained by estimating the correlation coefficients between all the frequency pairs in the output of the discrete Fourier Transform. Importantly, as discussed by Masimore et al. (2004; 2005), when implemented as an autocorrelation, this method allows the fundamental frequencies of the LFP to be identified without filtering as well as determine any potential interactions across different oscillatory bands. When implemented as a cross-correlation, it provides a measure of the degree in which the variance in power within one region affects the variance in power within another region.

### Ensemble empirical mode decomposition

The ensemble empirical mode decomposition (EEMD) analysis follows the procedure of Lopes-Dos-Santos et al.(2018). The EEMD was applied to decompose the original LFP into Intrinsic Mode Functions (IMFs). White noise was added to the original signal before decomposition to alleviate mode mixing, and was canceled out after the decomposition by averaging over ensemble (Wu and Huang, 2009). In this study, the ratio between the variance of added white noise and the original LFP was 0.5, and the ensemble number was 200. After applying the EEMD, the power spectrum for each IMF was used to identify the peak frequency. The supra-theta signal is defined as the sum of all the IMFs with peak frequency above 12 Hz. The time-scale power distribution of supra-theta signal was obtained by Morlet wavelet transform with constant ratio for the wavelet family set as 7 (Colgin and Moser, 2009).

#### III. Time-frequency atoms

For the Fourier transform, the duration *T* of the time sequence and the frequency resolution Δ*f* of the transformed sequence are related through the reciprocity relation

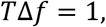

which implies that increasing the frequency resolution Δ*f* is equivalent to increasing the time duration *T* of the analyzed signal. The *T*Δ*f* = 1 equation highlights an important limitation of the Fourier transform: if *ɡ* (*t*) is highly localized, its transform *G*(*f*) has a wide frequency support. This means that very high sampling rates to cover the wide frequency domain, and the interpretation of the high-frequency content can become difficult. Restricting the Fourier analysis equation to a specified duration *T* is equivalent to multiplying the time series *ɡ*(*t*) by finite support rectangular window *w*(*t* − *τ*) = 1 if |*t* − *τ*| ≤ *T*/2 and zero otherwise. In other words, the integral operator

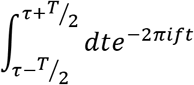

may be written as

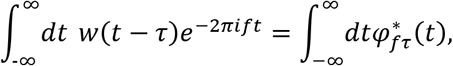

which may be interpreted as a projection of *ɡ* onto functions *φ*(*t*) = *w*(*t*)*ψ*(*t*). The function *φ* could be described as a localized oscillation. If one constructs in the time-frequency plane a rectangle of sides *T* and Δ*f* centered, say, at 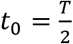 and 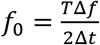, the *T*Δ*f* = 1 equation states that the area of this rectangle is constant, regardless of the value of *T*. This rectangle is sometimes called Heisenberg box (Mallat, 1999). This is in fact an example of the application of the general Heisenberg uncertainty principle, which states that the area of a Heisenberg box cannot be made arbitrarily small. For an arbitrarily-shaped localized oscillation *φ*(*t*) with Fourier transform *ϕ*(*f*) and unit variance 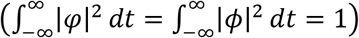, defining the time and frequency widths as

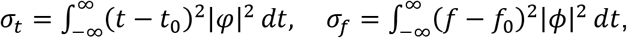

one can show (Gabor, 1946; Percival and Walden, 1993; Mallat, 1999) that 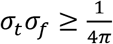.

In other words, it is impossible to achieve simultaneous arbitrary resolutions both in time and frequency. While the time-frequency resolution (area of Heisenberg boxes) cannot be made arbitrarily small, it can be minimized. Gabor (1946) showed that the minimal area for a Heisenberg box is achieved by localized oscillation *φ*(*t*) = *w*(*t*)*e*^2*πift*^ where *w* is a Gaussian-shaped window. Following his work, Goupillaud (1984) used this shape, later called the Morlet wavelet, to introduce the continuous wavelet transform (Grossmann and Morlet, 1984; Mallat, 1999). The limitations imposed by the Heisenberg uncertainty principle are illustrated in **figure 2**. A Morlet wavelet results as a product of a sine function with a Gaussian window. The resulting function is the “mother wavelet”. The wavelet transform then uses scaled versions of the mother wavelet as “elementary” functions.

**Figure 2:**
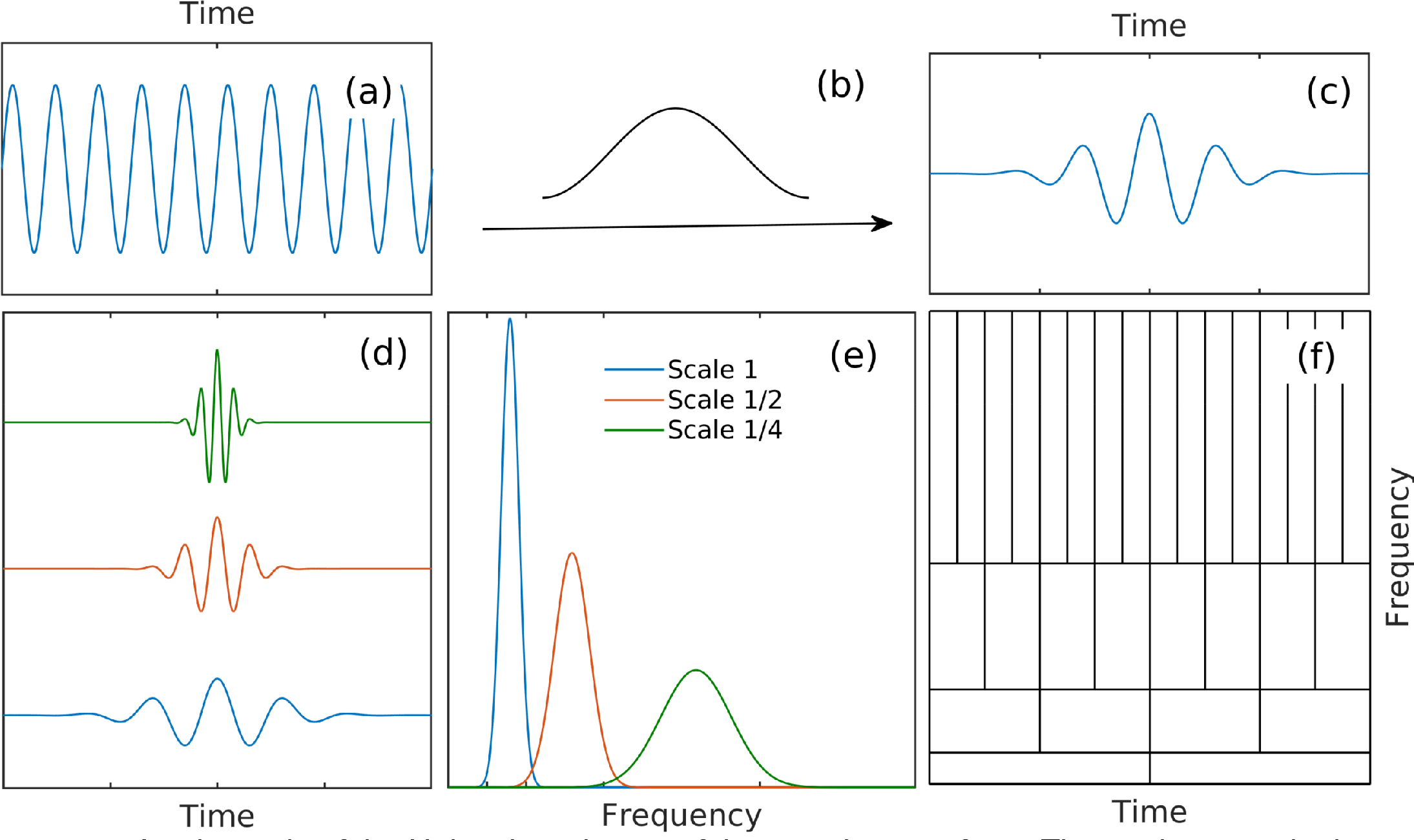
A schematic of the Heisenberg boxes of the wavelet transform. The mother wavelet is obtained by the multiplication of a sine function with a window (panels a-c). d) Wavelets at three scales resulting by the scaling of the mother wavelet. e) Frequency distribution of the power of wavelets shown in e). f) Heisenberg boxes of the wavelet transform.

It is important to note that the wavelets shown in figure **2**, panel d, are not harmonic, therefore they are *not uniquely characterized by a single frequency value*; instead, they are characterized by a frequency interval, say, corresponding to the width of the frequency spread of their power (as given by the Fourier transform). The frequency distribution of power is, as expected, narrower for the larger-scale wavelets, and wider for smaller-scale wavelets. Therefore, the wavelet Heisenberg boxes coverage of the frequency axis is not uniform, and consequently, using wavelets as a “frequency” representation results in a non-uniform frequency resolution, with resolution degrading at higher frequencies.

#### IV. Numerical implementation

Fourier and wavelet analysis procedures were implemented in the Matlab^®^ environment using available Matlab^®^ toolboxes. The Fourier cross-spectrum was estimated by dividing the LFP time series into 2048-point (1 second) segments sorted by rat speed and windowed using a hamming window; cross-spectra were estimated by averaging over all segments in each speed bin; the coherence and phase lag are defined as the modulus and the argument of cross-spectrum, respectively. The wavelet transform used the Morlet wavelet, with the central frequency of Morlet wavelet used as nominal frequency. The Morlet wavelet family is characterized by the constant ratio between the central frequency of wavelet and standard deviation of the applied Gaussian window, set to 7 in our analysis in order to match the methods of Colgin et al. (2009).

The wavelet scalogram is defined as the logarithm of the squared modulus |G_mn_|^2^ of the wavelet coefficients. The power spectral density for the wavelet transform were computed as the time marginal of the wavelet transform (Abry et al., 1993). The Fourier spectra were estimated using the Welch method (Welch, 1967).

### Methodological Limitation

At this point, it needs to be stressed that the wavelet and EEMD assignment of variance to a specific frequency (e.g., in the construction of a power spectral density) is ultimately arbitrary and potentially meaningless, since the elementary functions used for the decompositions <u>are not harmonics functions</u> (sine/cosine). The data presented here are for illustrative and replication purposes. We believe this methodological treatment is flawed and should be treated with circumspection

## RESULTS

The anatomical connectivity of the hippocampus is well conserved such that CA3 input into CA1 terminates into the str. radiatum layer of CA1 while entorhinal input terminates in both the oriens and str. lacunosum-moleculare (Sheremet et al., 2018b). As the entorhinal cortex is has been reported to convey fast gamma into the hippocampus whereas the radiatum supports slow gamma (Colgin et al., 2009), we first calculated the current source density LFP frequencies across rats during behavior. Figure 3 shows the current-source density plots for the theta frequency band as well as the first harmonic (15-17 Hz) and slow gamma (25-55Hz). With the evidence that prospective coding increases with running velocity (Maurer et al., 2012) and that prospective coding occurs during slow gamma epochs (Bieri et al., 2014; Zheng et al., 2016), we optimized for the detection of slow gamma by selecting epochs of rat speed greater than 35 cm/s. First, the current-source density of theta shows typical phase shift of theta rhythm from oriens to the dentate gyrus across rats, demonstrating a reliable reconstruction of electrode position (**Fig. 3**; also see Sheremet et al., 2018b). As the multiple gamma hypothesis contends that there are two if not more gamma generators in the hippocampus that support the high and low frequency oscillations (Colgin, 2015), we anticipated different source and sink patterns between the harmonic of theta and the gamma oscillations.

Unsurprisingly, the distribution of sources and sink for the theta and the first harmonic are nearly identical with both frequencies exhibiting maximum current deflection in the hippocampal fissure (compare with figure 2 of Bragin et al., 1995). However, of concern is the similarity of fissure sources and sinks between the traditional gamma range (25-55Hz) and both the fundamental and first theta harmonic. One sure way to cleanly differentiate oscillations would be through the demonstration of minimal synaptic overlap. These results, however, reveal that the synapses that terminate around the fissure support both theta, the harmonic and the 25-55Hz oscillation band (cf. figure 4C of Belluscio et al., 2012). Therefore, the current source density analysis is - at the least - inconclusive regarding the disambiguation of slow gamma from theta harmonics. At the worst, it suggests that precisely the same synapses contribute to the spectral frequencies between 8-55 Hz (suggesting that oscillations will inevitably be interdependent and coupled; evidence against the multiplexing hypothesis).

**Figure 3:**
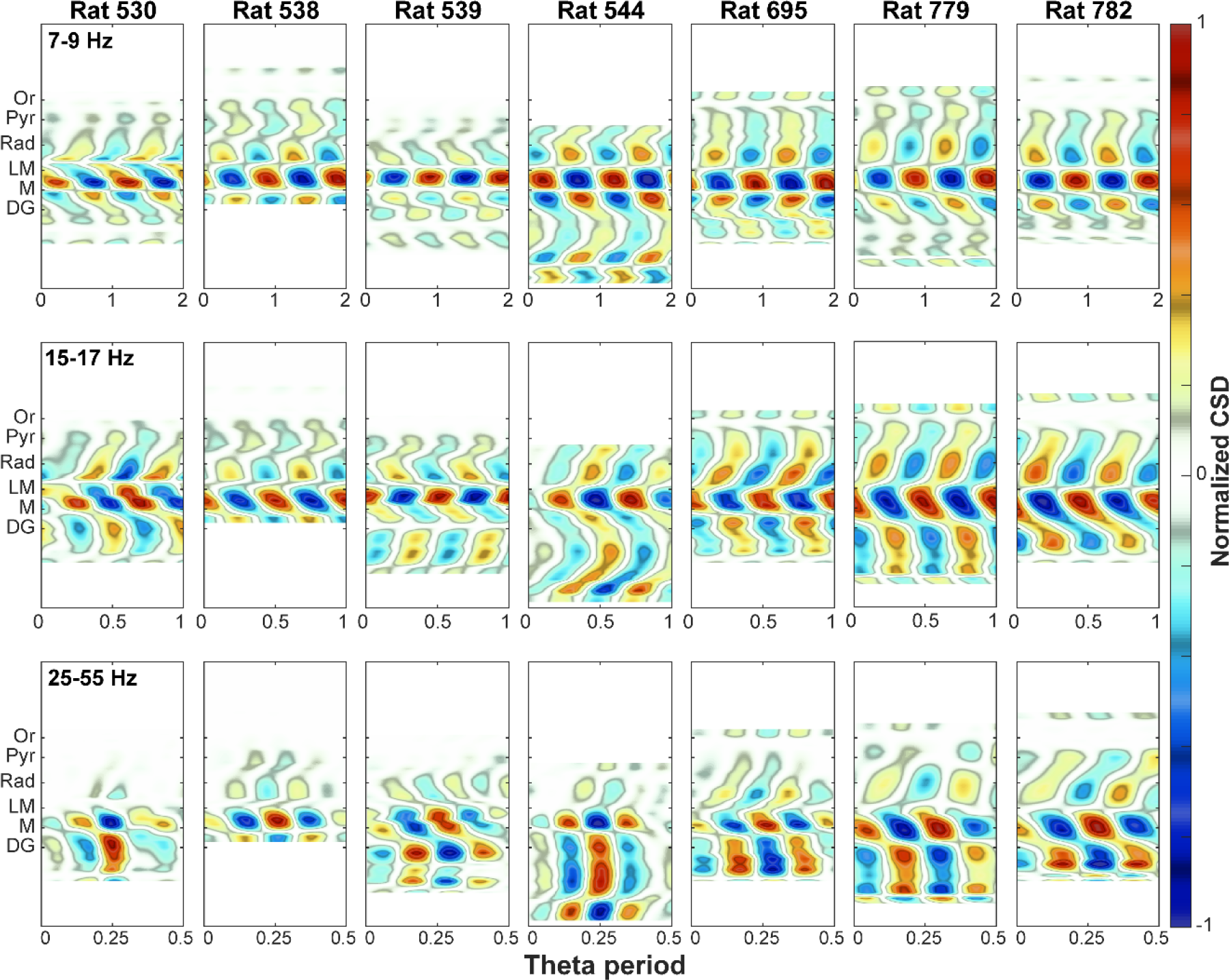
Current source density (CSD) analysis of seven rats during high speed movement. LFP recordings were split into 1 second segments and only segments with mean velocity larger than 35 cm/s were kept. LFP segments were aligned based on maximum theta peak in stratum LM and then were averaged. CSD analysis were carried out for three frequency bands: 1) theta, 7-9 Hz (top row); 2) theta harmonic, 15-17 Hz (middle row); 3) slow-gamma, 25-55 Hz (bottom row). For each panel, CSD values were normalized by maximum source/sink strength. In contour plots, warm color represents current source and cold color represents current sink. Considering the differences of probe positions and hippocampus sizes among rats, the CSD data were rescaled and shifted based on hippocampus lamina. In individual rat maps the vertical axis is channel number (not shown). Phase changes across channels (vertical) of the current-source density allow for the identification of oriens (Or), pyramidal layer (Pyr), stratum radiatum (Rad), lacunosum-moleculare (LM), molecular layer (M) and upper granule layer, or dentate gyrus (DG). It may be interpreted that the dentate is capable supporting a 25-55Hz frequency oscillation such as slow gamma (Colgin et al., 2009) or Beta (15-30 Hz; Rangel et al., 2015). We offer caution here as there are multiple theta diploes in the hippocampus with theta in the dentate potentially decoupled from theta in the str. radiatum (Montgomery et al., 2009). Furthermore, the theta oscillation in the dentate, cast a high number of harmonics relative to the other layers with the consequence of higher sources and sinks in the 25-55Hz band (see Fig. 5 below).

Comparison of these CSD results to the data of Belluscio et al. (2012; slow gamma defined as 30-50 Hz) and Lasztóczi and Klausberger (2016; slow gamma defined as 32-41 Hz) are partially at odds. Specifically, these studies identified a slow gamma rhythm in the radiatum. In particular, Belluscio and colleagues observed a significant current source density in the radiatum when the LFP was filtered in the slow gamma range (see Figure 4C of Belluscio et al., 2012). Therefore, we considered the possibility that due to unaccounted confounds, our results in Figure 3 were incorrect. In light of this, we analyzed a single shank of a 256-site silicon probe generously provided by the laboratory of Gyuri Buzsaki (Fernández-Ruiz et al., 2017). Current source density was calculated during ripples as well as the LFP during behavior filtered for theta, the first harmonic of theta, slow gamma (25-55 Hz) and a broad gamma (defined as 50- 120 Hz based on power spectral density cross-correlation and bicoherence analyses presented in Sheremet et al., 2018b). Similar to figure 3, we found nearly identical profiles between theta, the first harmonic, slow gamma and the broad gamma frequency (**Fig. 4**). Therefore, in yet another preparation, the radiatum has no more or less “slow gamma” than would be anticipated from comparing the power spectral density profile across hippocampal layers (Sheremet et al., 2018b). Furthermore, these results strongly parallel the original findings of Bragin et al. (1995). It should be noted that Lasztóczi and Klausberger (2016) implimented a wavelet analysis to identify different gamma bands with their slow gamma definition close to where the 32 and 40 Hz harmonics of theta would reside (Sheremet et al., 2016; Sheremet et al., 2018b). As the CSD failed to disambiguate oscillatory frequencies across hippocampal lamina, but prior reports using wavelet identified gamma, a new analytical course was determined.

**Figure 4:**
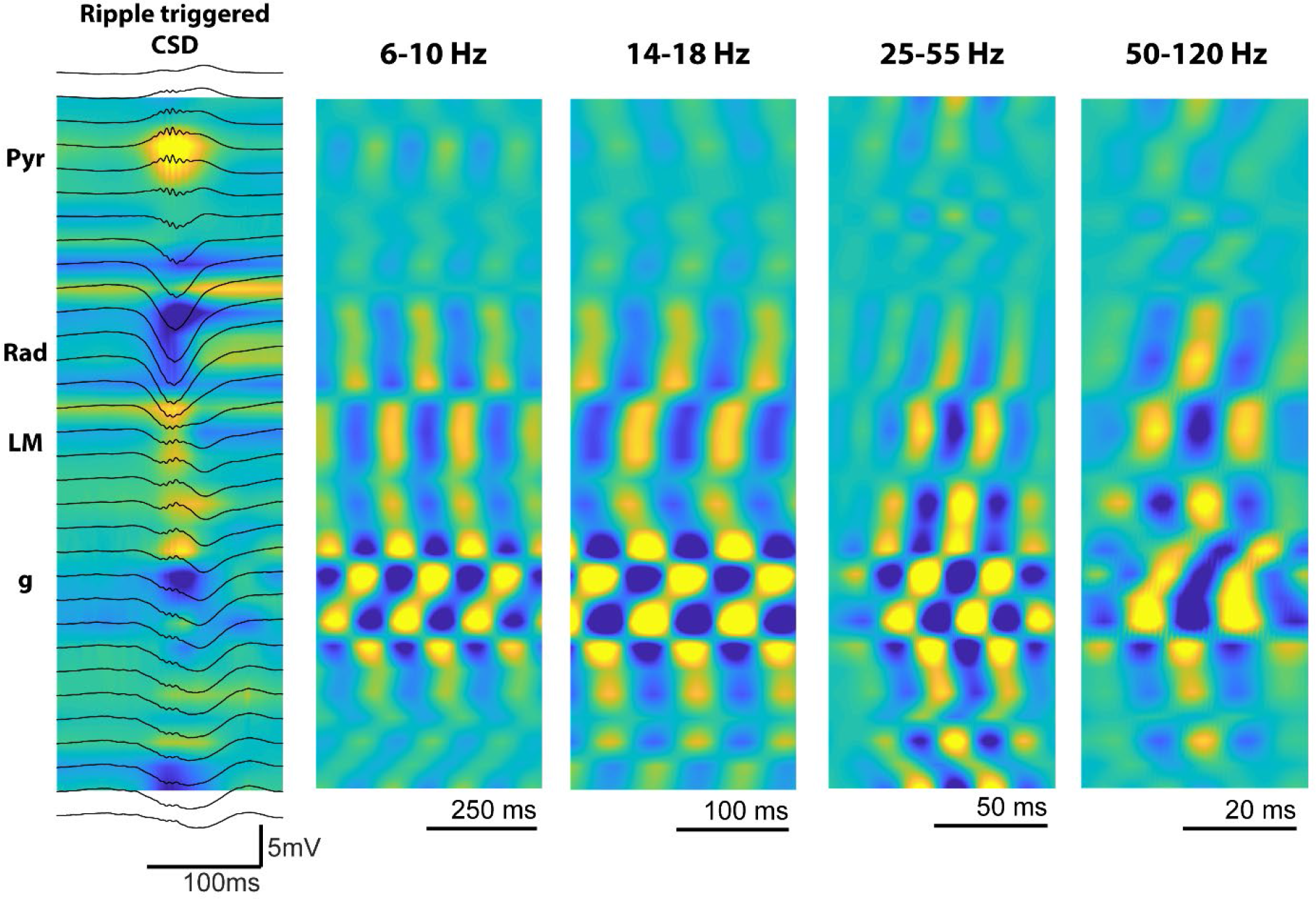
Current source density (CSD) from data generously provided by the Buzsaki laboratory (Fernández-Ruiz et al., 2017). The ripple triggered CSD clearly resolves sources and sinks associated with the pyramidal layer, radiatum, lacunosum moleulcare and dentate gyrus. The source sink profile is distinct between ripple events and theta. Note well, however, that CSD plots for theta, the 16 Hz harmonic, slow gamma, and traditional gamma, the source sink profiles are highly similar. Simply, current is generated where there are synaptic contacts. As there does not appear to be separate layers that support specific frequencies, a parsimonious conclusion is that the synapses themselves are agnostic to the LFP frequency that they support. (Pyr, pyramidal layer; Rad, radiatum; LM, lacunosum-moleculare; g, granule cell layer). Color axis scales: Ripple: −0.75 to 0.75 mV; Theta: −0.15 to 0.15 mV; 1^st^ Harmonic: −0.030 to 0.030 mV; Slow Gamma: −0.015 to 0.015 mV; traditional gamma: −0.01 to 0.01 mV.

Specifically, the overlap between the current sources of all oscilaltions works against the idea that there are distinct generators that decouples theta from gamma. Moreover, there seems to be limited rationale with respect to the sub-divion of the gamma band other than analytical methods of spectral decomposition. Earlier methods utilizing Fourier based decomposition only reported a unitary broad band gamma (Bragin et al., 1995; Buzsáki et al., 2003) whereas the initial identification of slow gamma implemented a wavelet decomposition (Colgin et al., 2009). As the Heisenberg boxes differ between the two methods of analysis (**Figs. 1** **and 2**), the wavelet “convolves” over the higher order harmonics (potentially accounting for why studies using wavelet decomposition rarely identify a harmonic above 16Hz). Therefore, as an initial benchmark comparison of Fourier decomposition versus a Wavelet decomposition, we generated a synthetic time series of pink noise embedded with an oscillation at 8 Hz and harmonics at 16, 24 and 32 Hz (**Fig. 5**). Notably the power estimate of the Fourier decomposition closely matched the theoretical distribution. However, as a consequence of the multi-resolution analysis of the wavelet decomposition, the power estimate significantly expands the frequency representation of the harmonics to cover a wide band, roughly overlapping with reports of beta (15-30 Hz; Rangel et al., 2015; Rangel et al., 2016) and slow gamma.

In addition to wavelet analysis, ensemble empirical mode decomposition (EEMD) has been put forward as a method theoretically capable of breaking the LFP into theta and supra-theta bands (Lopes-Dos-Santos et al., 2018). Specifically, Lopes-Dos-Santos and colleagues claim that, following EEMD, the remaining supra-theta signal is “harmonic free”. Therefore, we conducted a direct comparison of fast Fourier transform, Wavelet decomposition and EEMD from the lacunosum-moleculare of an awake-behaving rat as a function of velocity (**Fig. 6**). This comparison clearly shows that, as anticipated from figure 6, that both the wavelet and EEMD analytical approaches severely deviates from the Fourier representation at movement speeds greater than 15 cm/s. Notably, this results in an apparent “convolution” of the third and fourth harmonics of theta raising a significant cause for concern. In particular, wavelet and EEMD yielded results congruent with the idea of “low gamma” while under representing the harmonics of theta (which exist in the raw trace as a sawtooth shape; Buzsáki et al., 1985; Sheremet et al., 2016). This phenomenon occurs to an extreme in the dentate gyrus, which exhibits up to the 5^th^ harmonic of theta, spilling into the range that has been defined recently as beta (Rangel et al., 2015; Rangel et al., 2016) or potentially slow gamma (Hsiao et al., 2016).

**Figure 5:**
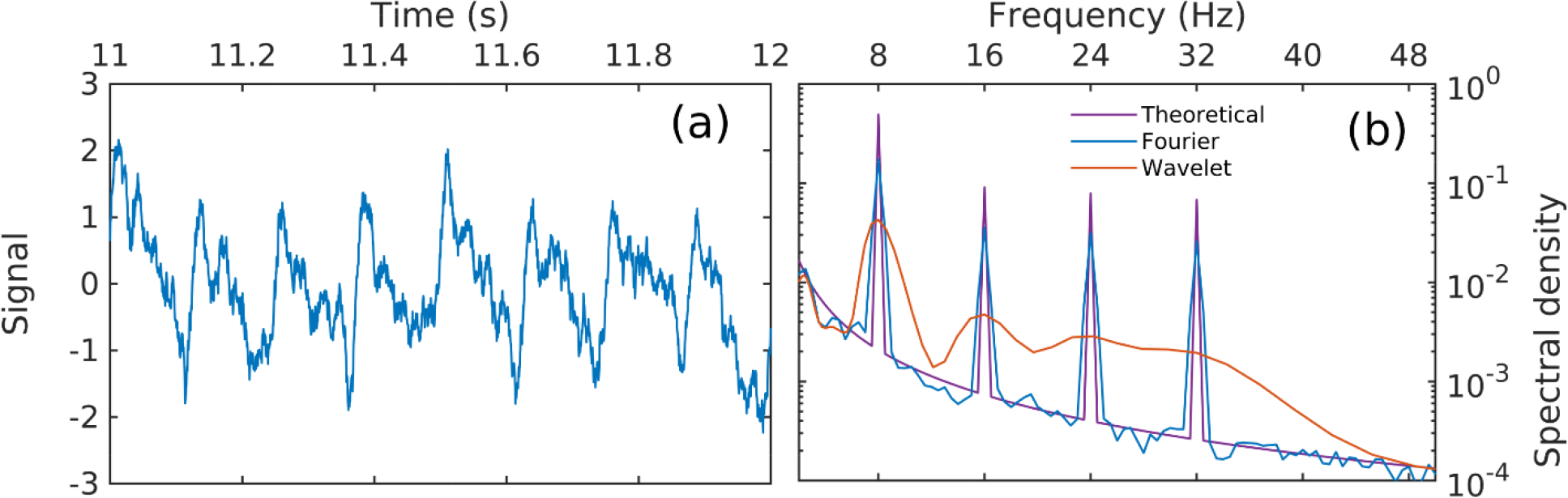
Example of Fourier and wavelet analysis of a synthetic time series emulating an LFP recording dominated by an asymmetric theta signal. The signal is constructed from an asymmetric sine function (an 8-Hz fundamental oscillation with phase-coupled harmonics), and pink (f ^−1.5^) noise. a) A time-series segment. b) Estimates of power spectral density.

**Figure 6:**
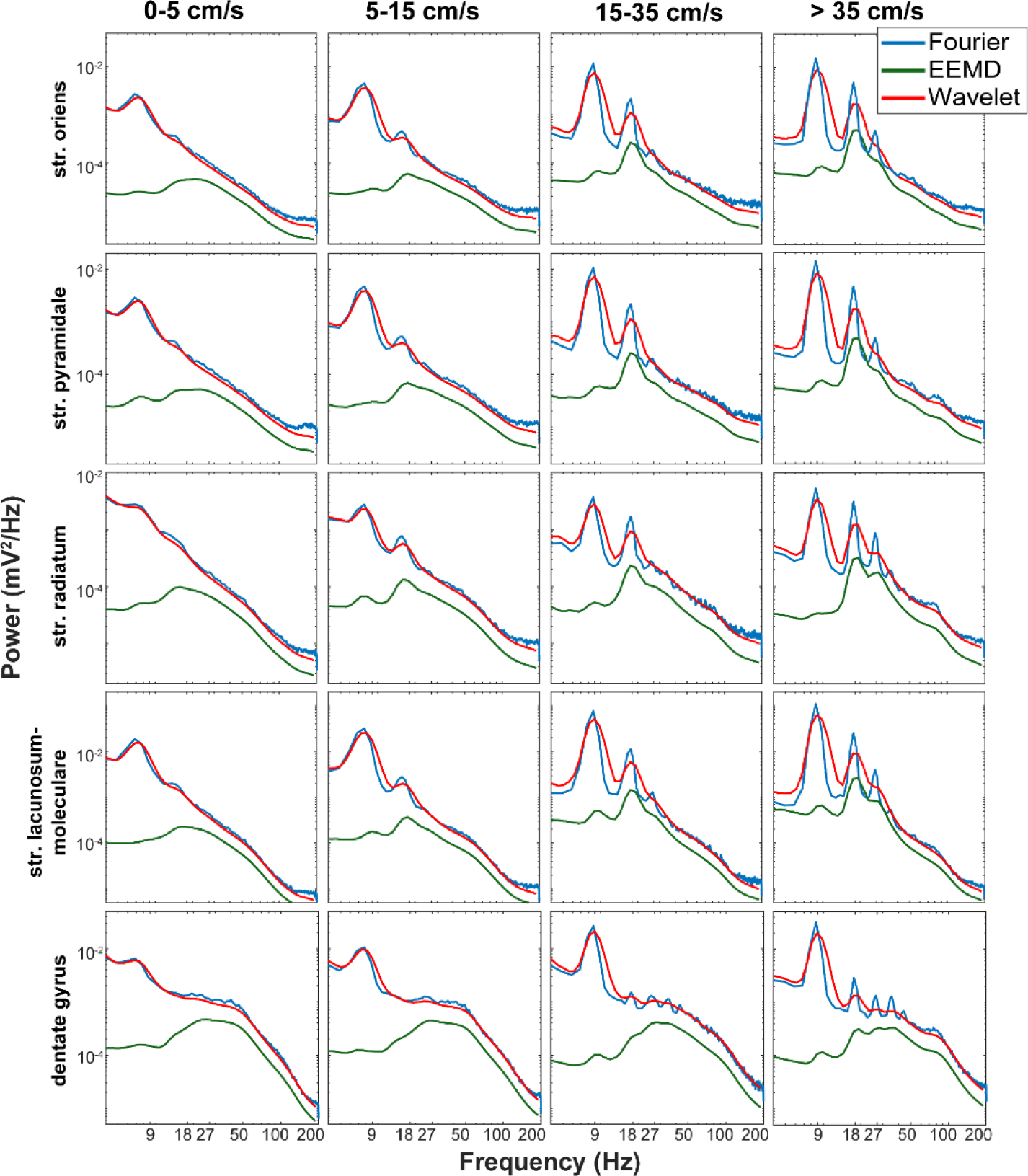
Comparison of three different spectral decomposition methods across layers and as a function of velocity. As previously describe by ourselves an others (Czurkó et al., 1999; Terrazas et al., 2005; Sheremet et al., 2016), Fourier decomposition clearly reveals the evolution of theta harmonics with velocity. This effect is consistent across layers, although the Fourier decomposition at high velocities in the str. radiatum and str. lacunosum-moleculare exhibits peak 32-36Hz range resembling a harmonic. However, wavelet decomposition based on the methods of Colgin et al., (2009) or the EEMD decomposition based on Lopes-Dos-Santos et al. (2018), due to the Heisenberg uncertainty boxes inherent in the analyses (Fig. 1) convolve over the 24Hz and higher order harmonics of theta. The peak of these broad bands roughly resembles what is described as “low gamma”. Also, please note that in the decomposition of the str. pyramidale, str. radiatum and dentate gyrus, wavelet analyses potentially underestimates the true power of the traditional 60-100Hz gamma (Fourier decomposition power exceeds the wavelet). This most-likely is a consequence of broad range averaging as the Heisenberg box becomes larger with higher frequencies. (Fig. 1). Finally, it Is worth noting that the removal of a fundamental oscillation, as in EEMD, reduces power in the theta rhythm, but fails to demonstrate the capability of addressing higher order harmonics. Rather, it parallels the outcome of the wavelet analysis. Once again, we remind the reader that, for wavelet and EEMD, *the assignment of variance to any specific frequency is ultimately arbitrary and meaningless*.

With this concern, we proceeded to revisit the wavelet scalogram analysis used by Colgin et al. (2009) to investigate the interaction between gamma frequency and the theta oscillation. This was first conducted for synthetically generated data (a sawtooth wave and an oscillation with integer locked harmonics) as well as data collected from the dentate gyrus and str. lacunosum-moleculare (**Fig. 7**). For the wavelet scalogram decomposition of the sawtooth wave, an infinite series of harmonics capture the instantaneous change in amplitude. The wavelet decomposition achieves this through a considerable peak in high frequency mid-trace. However, this is accompanied by multiple low-level harmonics within the slow gamma range. While this is a simulation to an extreme derivative, the time-frequency representation the synthetic harmonic series (8, 16, 24, and 32 Hz) is nearly identical to the slow gamma bumps that can be seen in the decomposition of either the dentate granule layer or the lacunosum-moleculare (compare Fig. 7 to figure 1Eb of Colgin et al., 2009). This “slow gamma” effect, however, can be accounted for the smoothing of harmonics observed in figure 6.

As the results of wavelet and EEMD are rarely portrayed as frequency versus power, but rather in terms of a time frequency representation, we furthered the comparison using two different windows of analysis (**Fig. 8**). Both in the long, three-minute fast Fourier transform and short, 6 second fast Fourier transform there is the clear presence of a 3^rd^ order theta harmonic (~24-27 Hz) and a trace of a 4^th^ harmonic (~32-36 Hz; **Fig.8**, **top panels**). However, as anticipated from the Heisenberg boxes (**Fig. 2**), examination of the wavelet on both the long and short time series result in a “smearing” and apparent intermittency of the ~24-27 Hz frequency band. Stated differently, the 24-27 Hz component is a direct consequence of theta being skewed and asymmetric (for example, more “saw tooth” shape than a sinusoid; Buzsaki et al., 1983; Buzsáki et al., 1985; Terrazas et al., 2005; Sheremet et al., 2016). This frequency is inherently coupled to the fundamental 8 Hz frequency, but the wavelet decomposition artificially detaches the 3^rd^ harmonic from the fundamental by using a different decimation of time and frequency (**Fig. 2**). This “bleed” and intermittency offers a strong resemblance to experiments that report different theta cycles exhibiting unique gamma signatures (e.g., Colgin et al., 2009; Bieri et al., 2014). To an extreme, using EEMD, it has been argued that theta can mix and match different gamma frequencies (Bagur and Benchenane, 2018) in support of different memory processes (Lopes-Dos-Santos et al., 2018). Therefore, we also replicated the methods of Lopes-Dos-Santos et al. (2018). Notably, comparing both the long- and short- temporal epochs of wavelet decomposition of the “supra-theta” reveals the distorted remnants of the second harmonic of theta as well as the third harmonic (**Fig. 8**). Finally, the summary power spectra for each method is presented demonstrating the inability of either wavelet or EEMD to clearly resolve the theta harmonics.

**Figure 7:**
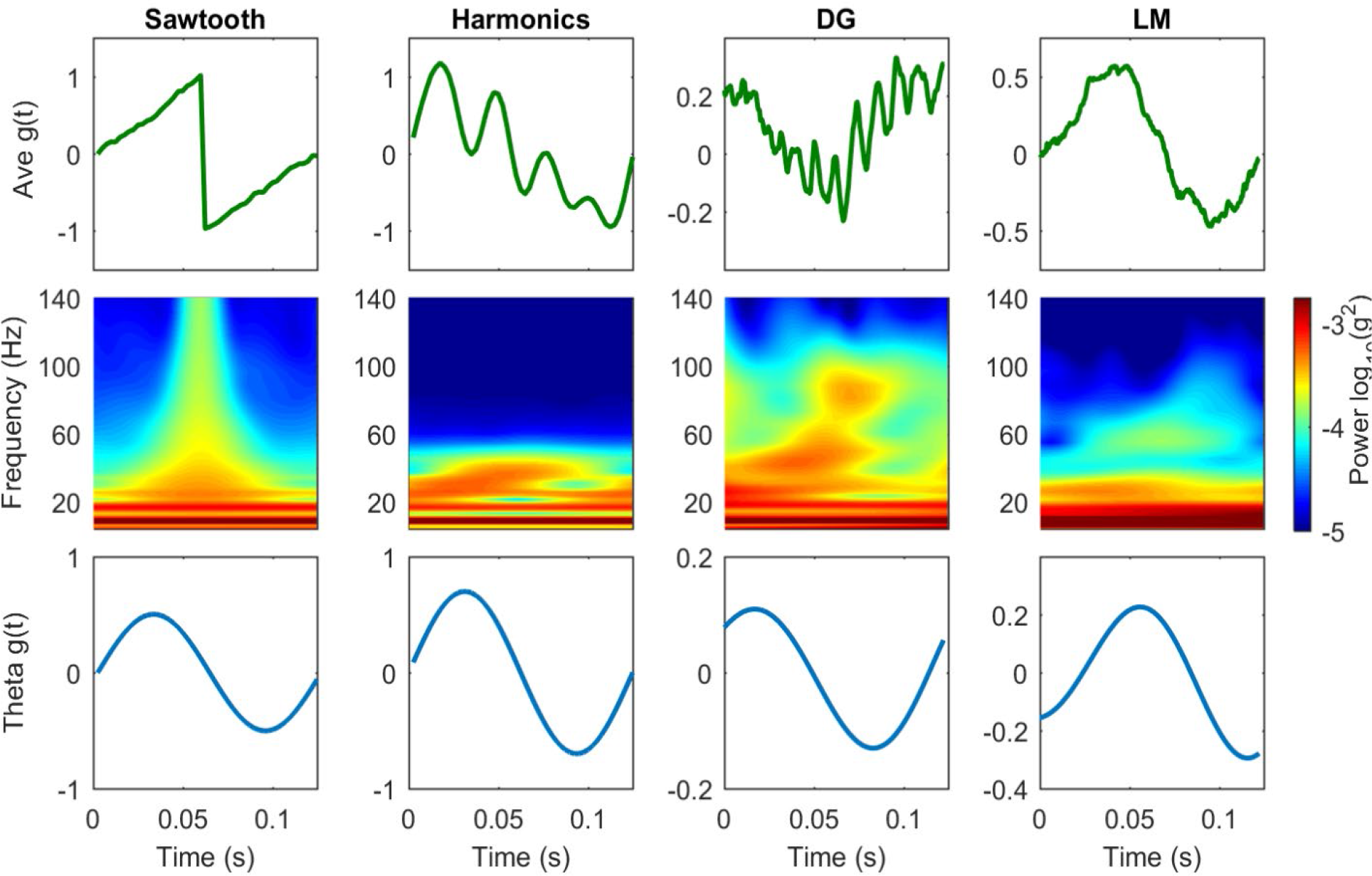
Wavelet scalograms for synthetic time series and LFP recordings in strata LM and DG. The averaged time series for one theta period (8 Hz) and wavelet scalograms were obtain for four time series: 1) Synthetic sawtooth waves (left to right, the 1st column); 2) Time series with fundamental 8 Hz oscillation and its harmonics (16, 24 and 32 Hz) (the 2nd column); 3) LFP recorded in DG (the 3rd column); 4) LFP recorded in LM (the 4th column). Synthetic time series have the total length of 10 seconds and were added with a background pink noise. LFP recordings were selected during high speed running with the total length of 5 seconds. The averaged time series were obtained by averaging over theta (8 Hz) periods (top row). The wavelet scalograms for averaged time series were computed with wavelet transform (middle row). Averaged time series were filtered within theta band (7.5-8.5 Hz) which represent the theta phase (bottom row). LFP data were collected from r530.

Up to this point, the current manuscript has focused on the primary methods of spectral decomposition that have led to the impression of a slow gamma oscillation in the hippocampus. However, this does not encompass the entirety of support for slow gamma. For example, analyses of hippocampal neuron spike times have revealed modulation to slow gamma (e.g., Colgin et al., 2009), with the logic being that as the cells exhibit a phase preference, one can sketch a theoretical mechanism by which they support the oscillation. The concern, however, has been raised that modulation in the 20-30Hz band may not actually be related to the slow gamma, but rather coupled to the asymmetry of theta (Schomburg et al., 2014). That is, the spike depth of modulation plots are not pure sinusoids, but exhibit skew and asymmetry (Skaggs et al., 1996; Quilichini et al., 2010). Furthermore, spectral decomposition of spike trains has revealed the presence an ~16 Hz harmonic modulation (Sheremet et al., 2016). Therefore, we considered the possibility that prior publications reporting phase coupling of neurons to slow gamma find significance for the reason that higher order harmonics in the LFP have a direct relationship to higher order harmonics in the frequency of spikes. Therefore, the point process spike trains (readily available on the CRCNS.org website; see Methods) were sorted into either a low or high velocity bin, converted into a binary timeseries, and either correlated against itself (autocorrelogram) or other neurons (cross-correlogram). These correlograms were then spectrally decomposed to examine the burst frequency modulation (see Geisler et al., 2007 for similar methods). Notably, in accordance with prior publications, the “power” (number of spikes) increases from low to high velocity and the burst frequency of the population increases with velocity (**Fig. 9**; Maurer et al., 2005; Geisler et al., 2007). Under the assumption that slow gamma modulation is highest when prospective coding is the greatest (Bieri et al., 2014; Zheng et al., 2016), which also describes the farther look-ahead at higher running speed (Maurer et al., 2012), it may be anticipated that there would be a single peak in the gamma range of the spike correlation spectrograms. Investigations of the spectrograms, however, do not support this perspective. Rather, the small observable peaks tend to coincide with harmonic lines (7.5, 15, 22.5, and 30 Hz). Therefore, the most reasonable conclusion is that phase coupling observed between a 25-55Hz band and spikes is carried by the harmonics evident in the LFP and the spike trains rather than being a consequence of a slow gamma oscillation that is distinct from theta. Note well that, when the power spectra are sorted by the frequency of the maximum power, pyramidal neurons (and their pairs) exhibit a greater frequency dispersion in the autocorrelograms and cross-correlograms relative to high velocity - *simply, oscillatory dynamics are stabilized by energy/input* (Churchland et al., 2010).

**Figure 8:**
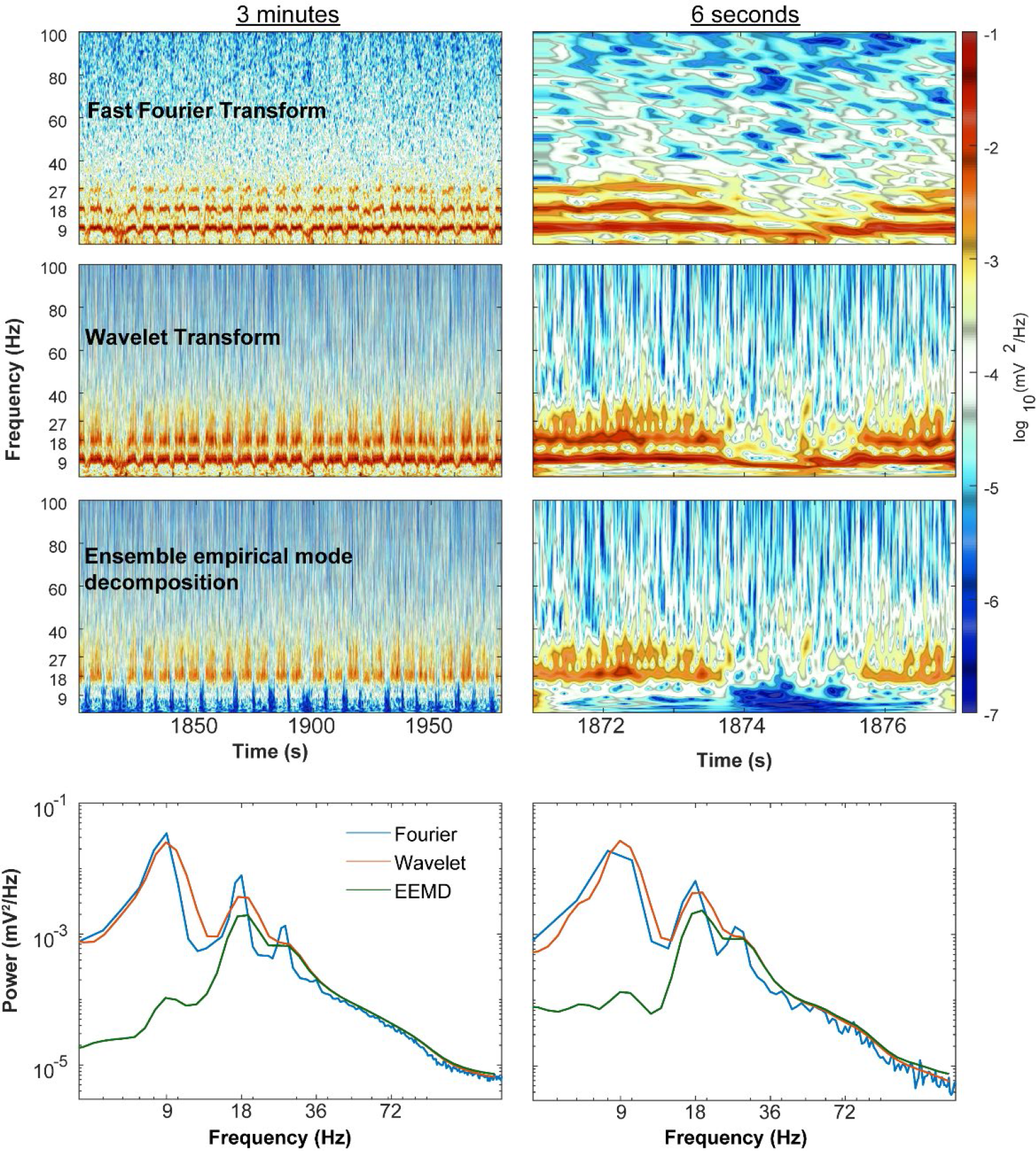
Spectrograms and power spectra during run epoch with different methods. Spectrogram and power spectrum were computed for a long run epoch (left column, 5 minutes) and a short epoch (right column, 6 seconds). For each epoch, the spectrograms were estimated with three methods: 1) Fourier transform (1^st^ row) with window length of 1 second and window increment of 0.1 second. 2) Wavelet transform (2^nd^ row) with Morlet wavelet. 3) Wavelet spectrogram of supra-theta signal (3^rd^ row). The supra-theta signal was obtained from ensemble empirical mode decomposition (EEMD) with noise level equaling to 0.5 total variance and ensemble number equaling to 200. The supra-theta was defined as the sum of decomposed modes whose central frequencies were large than 12 Hz. The wavelet spectrogram of supra-theta was computed with the method described above. Power spectra were computed by averaging the spectrogram over time (bottom row). For both epochs, the Fourier transform identifies theta and high order harmonics while wavelet analysis tends to resolve a theta rhythm and a wide-band frequency component (16-30 Hz). By considering this band as independent from theta in this manner gives the unintentional representation that there are “bursts” of gamma. Stated differently, the 24Hz oscillation is inherently dependent on the nonlinearity of theta (“saw tooth” shape of theta) and thus is incorrectly decomposed by the wavelet. Data were collected from r782.

Significant emphasis has been placed on coherence and cross-frequency analysis as a justification for slow gamma. Caution, however, has been emphasized with respect to low values of coherence (Buzsáki and Schomburg, 2015), alluding to the idea that low values have contracted meaning. Furthermore, under the hypothesis that hippocampal LFP patterns are organized by a cascade of energy from low frequencies to high frequencies (Buzsaki, 2006; Sheremet et al., 2018b), coherence should be significantly driven by activity into the hippocampus (which is approximately correlated with running speed). Therefore, we investigated coherence between the dentate gyrus and other lamina of the hippocampus as a function of velocity (**Fig. 10**). For comparison, we have also included the power spectral density measure and phase offset as a function of depth. At low velocities, coherence is fairly low without any notable deflections above theta. At high velocities, however, there is both an increase in the power of theta harmonics along with large coherence in theta harmonics across hippocampal lamina. Given the substantial increase in coherence across the theta harmonics with velocity, a traditional approach of considering coherence that does not account for velocity may unintentionally smear the range between 16-48 Hz, giving the impression of a single peak.

**Fig. 9:**
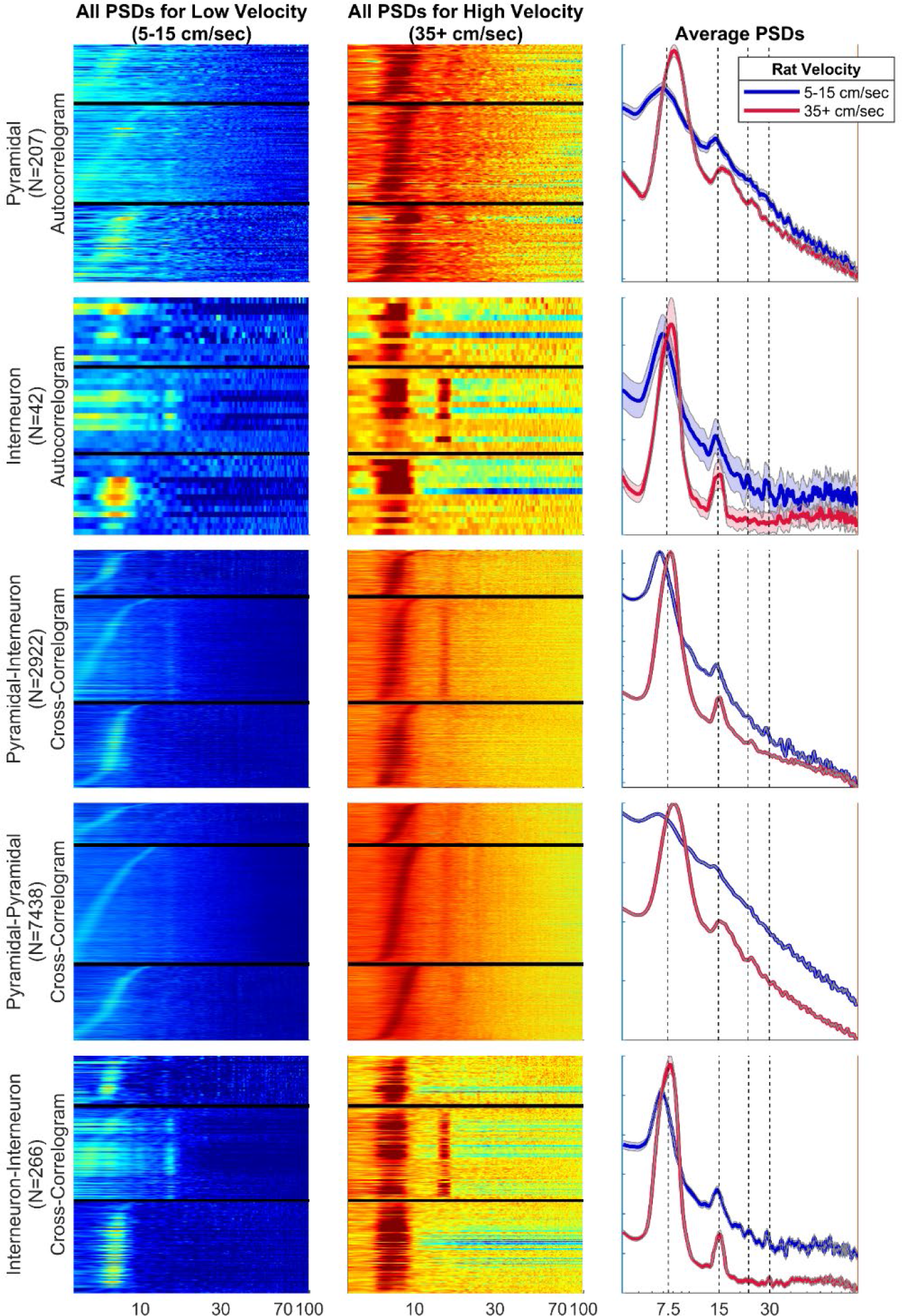
Power spectral density of neuron autocorrelograms (top two rows) or cross-correlograms (bottom three rows). The left column are the individual power spectra for auto- and cross-correlograms taken at slow velocity (5-15 cm/s) while the middle column are the same neurons or neuron pairs at high velocity (35+ cm/s; color axes are to the same scale between plot). The rows are sorted by the maximum frequency between 6-10Hz. The right column is the average power spectral density of the spike auto- and cross-correlograms at low (blue) and high velocities (red); errorbars are the standard error of the mean (different axes are used to allow spectral shape to be compared). The red dashed vertical lines are plotted at integers of 9 Hz up to 36 Hz. Note that neither the auto- or cross-correlogram power spectral show a single, slow gamma peak. Rather, should there be a notable deflection, it coincides with harmonics of the fundamental frequency. Also note that the dispersion in frequencies of the auto- and cross-correlations that include pyramidal neurons decreases as velocity increases. Energy/input stabilizes coherent oscillatory patterns.

Coherence analysis, however, to a large extent is unaffected by changes in power (of course, an amplitude must be present to define a phase). Therefore, we considered the argument that slow gamma is not entrained through the hippocampus in terms of phase-phase coupling. Rather, power is the important driver of coupling. Therefore, using the methods of Masimore et al. (2004; 2005), we investigated how changes in power within either the medial entorhinal cortex or dentate gyrus correlated with power changes across the lamina of CA1. In accordance with our recent publication (Sheremet et al., 2018b), we only found evidence of cross-regional power interactions between theta, theta harmonics and a unitary gamma (**Fig. 11**). From these power-power analyses, there is no reason to believe a slow gamma oscillation exists.

Finally, if the observation of slow gamma were a consequence of theta harmonics, then there should be a strong dependence on velocity. Investigation of the primary literature, however, is not so clear cut. For example, Chen and colleagues observed a linear change in slow gamma with velocity (2011), supporting the harmonic hypothesis. Ahmed and Mehta, on the other hand, found a reduction in slow gamma power with running speed (2012). Analytically, these results are not incorrect, but more nuanced. Specifically, as animal running velocity increases, there is an associated increase in theta power and the harmonics. These increases in power can be viewed as the energy into the hippocampus increasing resonance or bringing otherwise incoherent neural activity into coherent, dynamic patterns. The consequence is that the incoherent activity surrounding the theta and harmonic bands: 1-6, 10-14, 19-23, and 25-40 Hz will effectively lose power (**Fig. 12**). Note well, this idea is against the idea of spectral multiplexing, in which each frequency has an independent meaning. Thus, the parsimonious explanation is that neurons can “drift” (exhibit a range of oscillations) when provided a low level of input, but become strongly entrained with energy and velocity (Sheremet et al., 2018b). This accretion of power in one or multiple bands comes at the expense of losing power in others.

**Figure 10:**
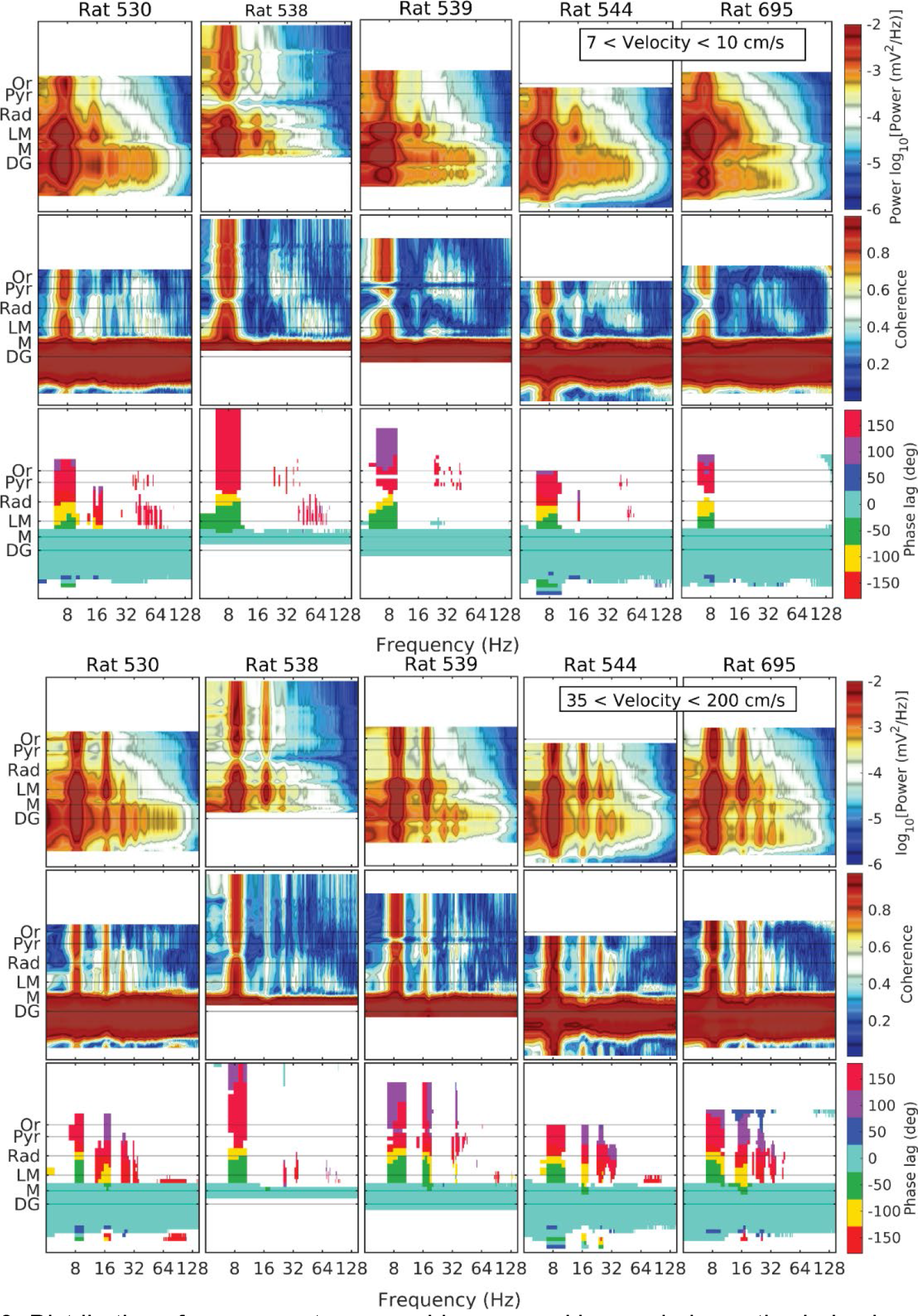
Distribution of cross-spectrum over hippocampal layers during active behavior, at low speed (top three rows) or high speed (bottom three rows). Cross-spectra shown are estimated with reference to the DG layer. For each rat, the observed regions of the hippocampus Top: spectral density of LFP variance. Middle: coherence (cross-spectrum modulus). Bottom: phase lag (cross-spectrum argument). On each sub-panel, the vertical axis is channel number (not shown); the panels are aligned according to the layer identification procedure. Distribution of cross-spectrum over hippocampal layers during active behavior), at high speed. Note that at high velocities, the power spectra, coherence and phase measures support the observation of multiple theta harmonics.

Nevertheless, this effect of spectral reorganization - eroding background with energy-likely accounts for the results also observed in Kemere et al. (2013) and Zheng et al. (2015). If anything, as oscillations are often discussed through the lens of entrainment, the disappearance of slow gamma power with energy/velocity supports the idea that it most-likely was never an oscillation at all, but rather a component of the noise spectrum.

**Figure 11:**
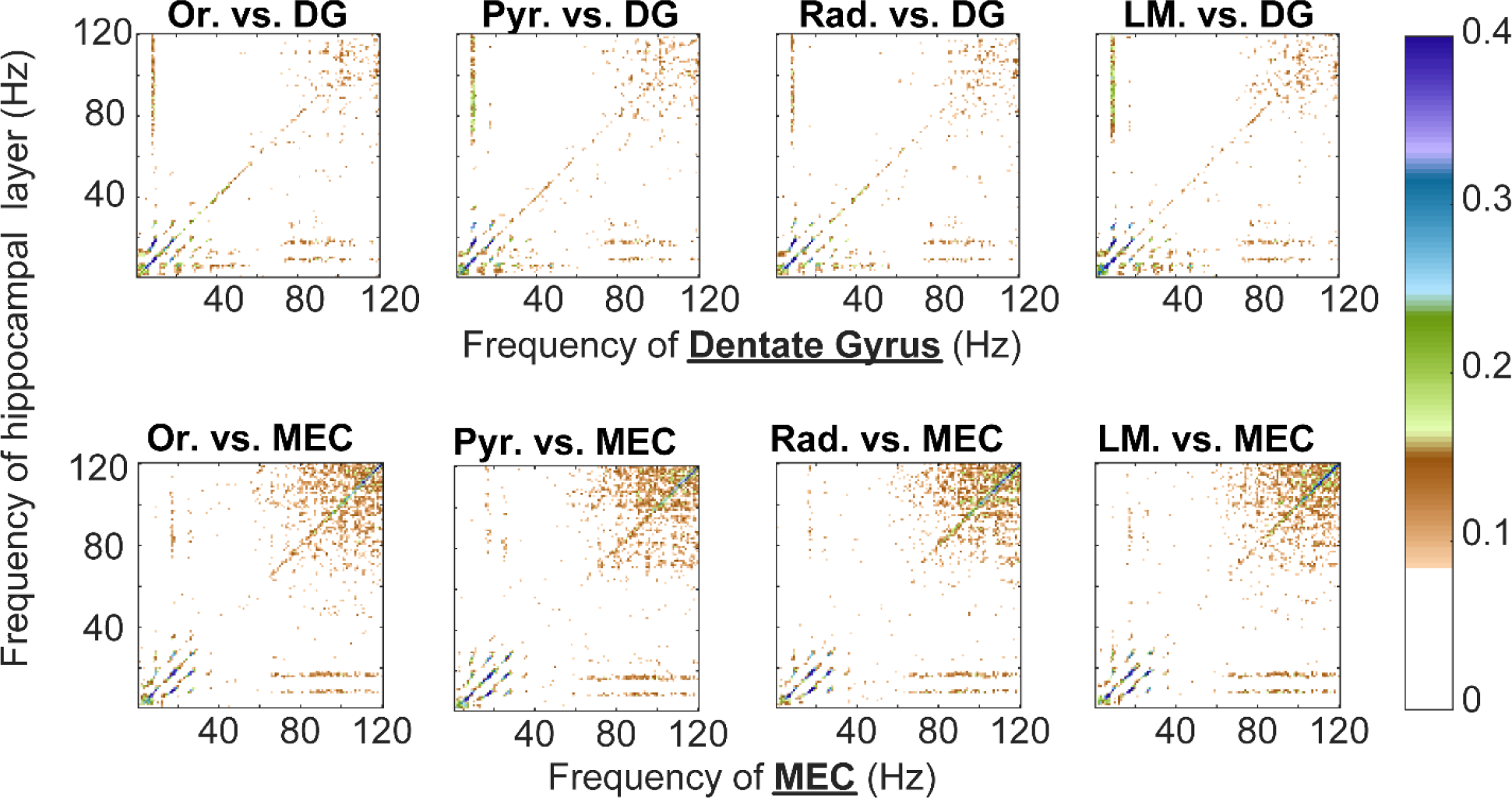
Average cross correlation coefficients of Fourier transform across three rats with electrodes in the medial entorhinal cortex in order to identify fundamental frequencies of interaction. As previously described, regions of non-zero correlation potentially indicate that two oscillatory bands showed high power at similar times suggestive of an interaction (see Masimore et al., 2004; 2005). In both cross-correlations between the DG and hippocampal lamina as well as the MEC and hippocampal lamina, there are notable “dots” of correlations below at ~40Hz and below, indicative of theta and its harmonics (Sheremet et al., 2016; 2018). Further off-axis, there is a correlation between ~80-120 Hz and theta as well as its harmonic. Notably, there is no reason to expect the need to subdivide gamma into multiple bands. Rather, the interactions are constrained to theta, its harmonics and the broad gamma band as originally described by Bragin et al. (1995; 50-120 Hz). For all plots, values below 0.03 are set to white. Or, oriens; Pyr, CA1 pyramidal layer; Rad, Radiatum; and LM, lacunosum-moleculare.

**Figure 12:**
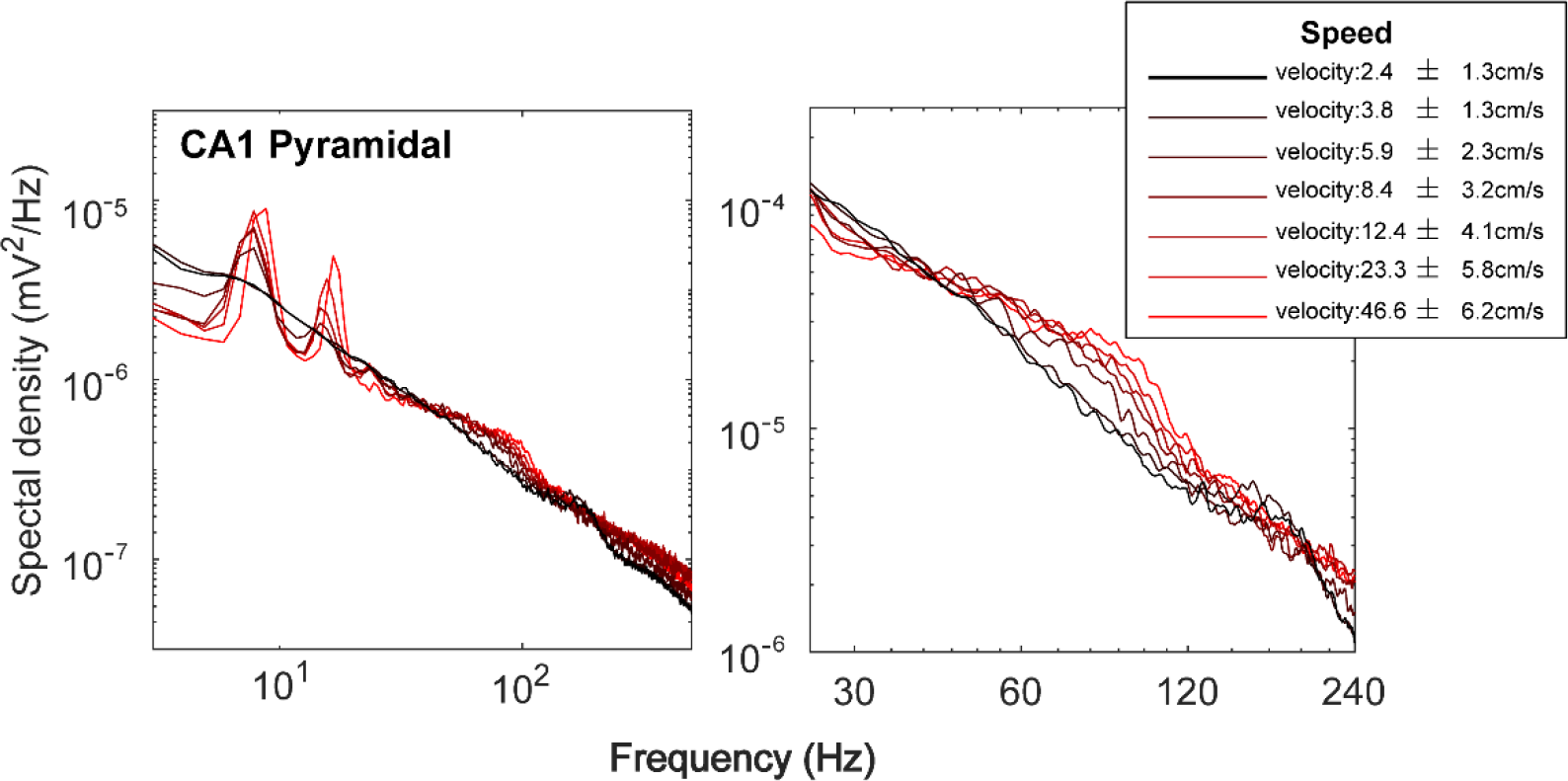
Spectral changes with velocity. It is often reported that there is a either no change or a decrease in slow gamma with increasing running velocity. While this is analytically sound, it is most-likely a result of the same reduction in power from 1-6, 10-14, 19-23, and 25-40Hz. That is, when a network of neurons receives a low level of input, the activity is unorganized, resulting in a smooth decrease in power from 1- 100Hz. However, as velocity is associated with increasing firing rates, the network organizes as a function of increasing energy (synaptic activity). Input stabilizes oscillatory dynamics (see Fig. 9; for similar data, see Churchland et al., 2010). By acting on one another, neurons will self-organize into a pattern of activity that supports theta, its harmonics and gamma. Thus, the reduction of the 25-40 Hz band is most likely a consequence of neurons becoming entrained. This effectively challenges the argument for slow gamma (it may be considered inappropriate to suggest an oscillatory band disappears when neural entrainment occurs). Compare to Ahmed and Mehta (2012; Fig 2).

## Discussion

The identification of slow gamma was initially predicated on wavelet decomposition (Colgin et al., 2009). More recently, empirical mode decomposition (EEMD) has been reported to find multiple low-frequency oscillatory bursts (22 Hz beta, 35 Hz slow gamma, and a 54 Hz mid-gamma; Lopes-Dos-Santos et al., 2018). Using the methods of Colgin et al. (2009) and Lopes-Dos-Santos et al. (2018), we successfully replicate these findings, but the results are contradicted by Fourier analysis. Specifically, the Fourier decomposition exhibits significant peaks in the high-order theta harmonics (24, 32, and 40 Hz). Critically, these contradictory outcomes are incompatible, necessitating a reevaluation of multiple concepts.

The fundamental issue is that a consistent, *physically meaningful* definition of the “rhythm” concept is missing, leaving one to fill the space with whatever seems handy. This defaults to the crudest of methods: visual inspection of the power distribution across frequencies. If the distribution is “flat”, that is, “does not have any eye-catching details”, it is usually called “pink noise” and assumed to contain nothing of interest. If the distribution has some evident details, say local peaks, then the corresponding frequency bands are declared worthy of attention and sometimes receive the “rhythm” designation. An example of this approach is the theta rhythm, which corresponds to a local maximum of significant magnitude, located around 8 Hz (Vanderwolf and Heron, 1964; Vanderwolf, 1969). Because the power in the 6-8 Hz band stands out, we consider it reasonable to assume that it has a special meaning. Distinguishing frequencies based on their power seems peculiar, since in fact any frequency band with non-zero power is in the Fourier representation an oscillation.

Note that, defined this way, “pink noise” and “rhythm” are arbitrary taxonomy concepts with no physics support. The obvious problem with this approach is that it leaves the door open for abuse. For example, to identify the gamma rhythm, one needs to generalize the ad-hoc “peak above noise” definition of a rhythm to something even more subtle, and perhaps less justified: in all cases but high running speed, there is no “local maximum” in the power spectral density in the gamma frequency band. Rather, the gamma band exhibits only an accumulation of power, with the spectral distribution remaining monotonically decreasing, but at a slower rate. This is a deviation from the flat, “pink noise” that is often made evident by compensating (e.g., whitening) the spectrum. However, finding peaks after standard whitening procedures are problematic if the spectra does not follow a unique slope, as is generally the case for hippocampal LFP (Sheremet et al., 2018b; Sheremet et al., 2018a). This raises the question “what is so special about spectral deviations that single out those frequency bands from ‘noise bands’ that have comparable energy?”. The answer seems to be the flat, pink noise spectrum has a different meaning and does not really matter enough to be called a rhythm. Only accumulations of energy past this background level are significant. The significance of these accumulations of energy (or losses of energy as seen in the 25-40 Hz band with velocity; Fig. 12), however, is unclear in the absence of a physical model of the phenomenon. Such a model should explain both the processes underlying the “non-rhythm” background spectrum, and the “rhythm” spectral deviations (e.g.,Sheremet et al., 2018a). In failing to do so, the multiplexing model of cognition seems improvised to account for the wavelet decomposition results. One must conclude that the definition of a “rhythm” based on the power distribution is an arbitrary decision, because it is based on the “aspect” of subjective representations and not on a fundamental, analysis-invariant description of the physics underlying the LFP process.

In contrast to the multiplexed “spectral finger print” perspective (e.g., Canolty et al., 2010; Siegel et al., 2012; Knight and Eichenbaum, 2013; Watrous et al., 2013; Akam and Kullmann, 2014; McLelland and VanRullen, 2016), the perspective that the hippocampal LFP is generated by the superposition of electrical fields generated by microscopic elements of a large, macroscopic system (Buzsáki et al., 2012) suggests statistical mechanics as a general framework for understanding LFP characteristics. While transient behavior in large nonlinear systems is possible (e.g. the sharp-wave/ripple complex), the behavior we expect for a “stable” system is the weakly nonlinear type, characterized by stationarity. In such systems the Fourier representation remains valid and the power spectral representation is defined. An alternative description of LFP processes, based on statistical mechanics and seeking to account for all processes expressed in the LFP spectral distribution (“rhythm” or not) is proposed in Sheremet et al. (2018a).

Therefore, when discussing the meaning of the “slow gamma rhythm”, it is important to examine carefully the assumptions about the underlying physical process embedded in the methods of power-frequency decomposition. Specifically, the observation of novel oscillatory bands have been supported through wavelet analyses (e.g., Colgin et al., 2009), and ensemble empirical mode decomposition (EEMD; Lopes-Dos-Santos et al., 2018). The issue of analytical applicability of each method to the hippocampal LFP necessitates a discussion on whether or not the experimenter considers the trace to be stationary or not (See Appendix). Nevertheless, as emphasized in the methods, wavelet and the EEMD approach of assigning power to a frequency is ultimately arbitrary and potentially meaningless, since the elementary functions used for the decomposition are not harmonic (sine/cosine). In other words, the power to frequency assignment becomes obscured.

Other empirical support for the existence of slow gamma included the correlation of units to the LFP (Colgin et al., 2009; Schomburg et al., 2014; Fernández-Ruiz et al., 2017), within and across region cross-frequency coherence (e.g., Colgin et al., 2009; Kemere et al., 2013; Hsiao et al., 2016), and cross-frequency coupling (e.g., Colgin et al., 2009; Belluscio et al., 2012). While these are supporting pieces of data, they neither confirm or deny the presence of an oscillation. Nevertheless, we have addressed each of these in the current manuscript. Notably, rather than perform a second-order analysis of investigating either phase locking to LFP or spike-LFP coherence, we simply sought to determine what frequencies are evident in the action potentials, using Thomson multitaper decomposition on auto- and cross-correlograms (Leung and Buzsáki, 1983). We did not observe any slow gamma modulation, but did observe multiple harmonics of theta (**Fig. 9;** Sheremet et al., 2016). Therefore, a wide band filter of 25-55Hz will most-likely result in a significant phase modulation of spike firing (e.g., Colgin et al., 2009; Schomburg et al., 2014) because-as noted by Schomburg and colleagues (2014)- this region covers the harmonic range. As pyramidal cells spike at theta frequency (and the first harmonic) and the harmonics can extend as high as 48 Hz (Sheremet et al., 2016), then the broad filter of 25-55Hz will be harmonic dominated. Furthermore, we have replicated cross-regional coherence, this time using velocity as a parameter, to investigate whether it is tenable to believe a broad slow gamma band exists. Although prior research would suggest that the dentate entrains other regions through slow gamma (Hsiao et al., 2016), our analyses here (**Fig. 10**) and elsewhere (Sheremet et al., 2018b) reveal coherence up to the 40Hz harmonic of theta. Furthermore, cross-correlation of power between the medial entorhinal cortex or dentate gyrus and layers of CA1 fail to show any interactions between frequencies other than theta, its harmonics, and a single 50-120 Hz gamma band (**Fig. 11**). Finally, we have addressed all forms of cross-frequency coupling - phase-phase, phase-amplitude, and amplitude-amplitude-in another manuscript (Sheremet et al., 2018b). Therefore, when considering both the primary and secondary evidence supporting the existence of slow gamma, there appear to be fundamental, critical shortcomings.

Of course, our observations do not cover the compendium of manuscripts that have supported the existence of slow gamma. An attempt to refute them all may prove to be a Sisyphean task. It is, however, worth noting that independent component analysis has been suggested as another method that reveals slow gamma (see Supplemental methods of Schomburg et al., 2014). Independent component analysis takes advantage of the conserved anatomy in order to identify source specific, non-volume conducted components localized to the layer. Notably, the range of slow gamma in Schomburg et al. (2014), 30-80 Hz (also used in Fernández-Ruiz et al., 2017), significantly differed relative to the prior publications of Colgin et al. (2009), 25-55Hz, and Belluscio et al. (2012), 30-50 Hz. The extended range of slow gamma in these papers were based off of a gamma amplitude-theta frequency (GA-TF) comodulogram and buttressed by the wavelet based Gamma amplitude - theta phase (GA-TP) modulation plots (see figures 2 and 3 of Fernández-Ruiz et al., 2017). Although the correlation coefficients of the spectrogram presented in the current manuscript (also see Sheremet et al., 2018b) stand in contrast to the comodulogram of Schomburg et al. (2014), we cannot speak directly to these results as we did not present data implementing their analytical approach. We can only suggest caution in running a preprocessing algorithm that involves applying principle component analysis and whitening prior to independent component analysis (see Supplemental methods of Schomburg et al., 2014) as it moves further away from the collected data.

To our knowledge, all of the aforementioned spectral decompositions and secondary analyses that report slow gamma fail to account for harmonics as suggested by Aru and colleagues (Aru et al., 2014). Using bicoherence to control for harmonics reveals that the low end of the spectrum is dominated by integers of theta, phase locked to the fundamental up to 48 Hz (Sheremet et al., 2016; Sheremet et al., 2018b; Sheremet et al., 2018a). From this analysis, there is no reason to believe that an independent slow gamma oscillation exists in the normal rat. However, our failure to observe this oscillation does not mean that the brain cannot venture into a parameter space in which slow gamma is prominently detectable from noise. It is therefore worth discussing what empirical data would be needed to support the assertion of slow gamma. Control for theta harmonics using bicoherence analysis should be required. As the argument has been made that slow gamma is coupled to theta, a strong interaction in the bicoherence between slow gamma and theta would further enhance this claim. Notably, the bicoherence analysis detects a strong coupling of the 50-120 Hz gamma to both theta and its first harmonic (Sheremet et al., 2018b). It should be mentioned that “wavelet bicoherence” analysis (Van Milligen et al., 1995) would suffer similar pitfalls to wavelet time-frequency representation. Moreover, considering asymmetry alone (e.g., Amemiya and Redish, 2018) is an insufficient control as oscillatory skew can also contribute to harmonics (Sheremet et al., 2016). Furthermore, as discussed above, oscillatory identification is partially determined by a deflection that significantly deviates from the “background” activity. Therefore, a simple, unwhitened, log-log power spectral density with a peak in the 25-55 Hz band using the Fourier transform would be indicative that there is a uniform signal that exceeds noise. Finally, controlling for animal velocity when conducting these analyses should become common practice. The secondary analyses should only be implemented once due diligence has been employed to ensure that a signal exists.

Through this presentation, we believe that fundamental, critical analytical inconsistencies are responsible for “slow gamma” detection that are instead a consequence of the theta harmonics. Moving forward, vigilance is necessary to ensure that the analytical treatment is “true to the essence of the biological process studied” or otherwise risk generating mistakes as a consequence of inappropriate methods (Marder, 2015). While quantitative data analysis reflecting what can be observed qualitatively is the first step toward a model, caution should be taken prior to applying analytical techniques that incorporate specific assumptions about the system without a physical model.

## Appendix Fourier and wavelet representation of deterministic time series

As mathematical, abstract transforms of functions defined on the real axis, both the Fourier and wavelet representations work fine, in the sense that they reconstruct the signal exactly (have inverses). In terms of applications to observational data, however, the two transforms were designed to meet different efficiency criteria.

The Fourier and wavelet analysis function on similar principles, decomposing the signal into “elementary” functions. However, the similarity ends here. In the Fourier transform case, the “elementary” are harmonic functions uniquely defined by their frequency; they form an orthogonal basis; hence the representation is unique and has a number of other convenient properties (e.g., conserves variance). In the wavelet transform case the “elementary” functions (wavelets)are a two-parameter (scale and position on the time axis) family of copies of a fixed shape, the mother wavelet. In contrast to the harmonic Fourier modes 1) wavelets are only required to have near-zero mean and be integrable in some way (fast enough decay at infinity) but are otherwise arbitrary; 2) wavelets do not form (in general) a basis (the representation is in general not unique); and 3) the result is a time-scale representation.

While the Fourier transform exists for a wide class of reasonably behaved functions and may be extended generalized functions. Its usefulness is illustrated by the Heisenberg uncertainty principle (see **Figs, 1** **and 2**) which implies that a highly localized signal has a wide Fourier spectrum. For example, if the time signal is a narrow pulse, the Fourier representation will require many high-frequency components to construct the sharp time variation - hence the wide spectrum. This also implies that the interpretation of the spectrum in this case is difficult, because the frequency components are phase locked to create the time pulse, hence they are not independent oscillations. Therefore, the Fourier representation is best applied to wide-support time functions, i.e., functions having some sort of translational near-invariance in time will have (we shall call this provisionally “stationarity”).

The wavelet transform analyzes signals by shifting in time bits of oscillations with different scales (**see** **fig. 2**). This approach is clearly efficient for signals with intermittent bursts of energy. For stationary (in the sense used above) signals, wavelets are inefficient because they use a two-parameter space when only one is really needed (see the Fourier decomposition). The decomposition also loses its intended meaning of identifying intermittent bursts.

Other than paying attention to these elementary efficiencies, the choice of representation (Fourier vs wavelet) for a given time series is arbitrary. There is, however, an elementary fallacy of reasoning that is often encountered: while they are both equally applicable, the results are not interchangeable. We stress that the wavelet ordering parameter is scale (the value of the dilation parameter) and not frequency. While scales are often translated to frequencies for ease of interpretation, one should not be confused: Wavelet “frequencies” are, strictly speaking, just labels. Wavelet have a non-trivial frequency distribution of power, i.e., “comprise” an entire range of (true) Fourier frequencies. Referring to them as if they had the Fourier meaning is incorrect and may lead to erroneous conclusions.

## Fourier and wavelet representations of stochastic processes

The concept of a stochastic process assumes that such time series are in fact random “realizations” of a “virtual” process, and the goal of analyzing the realizations is to characterize this virtual process, and not its individual realizations. For example, averaging a quantity characterizing the stochastic process is a procedure that takes all possible realizations and computes a representative value that characterizes the stochastic process and not individual realizations. Such averaged quantities are called in practice “estimators”, exactly because they are assumed to converge to a well-defined value if the averaging is done over the entire realization space. In practical applications, the stochastic process whose realizations (time series) are observed is not entirely accessible, therefore the existence of a particular average quantity for an arbitrary process. This implies that one does not have a rigorous characterization of the properties of the stochastic process. This is, however, essential for interpreting the observations and insuring that one does not elaborate on meaningless, at worst non-existent, quantities. It is important to understand that just because a finite number of realizations always yields an average does not necessarily imply that the stochastic average (over all realizations) exists. Therefore, in practice, one is forced to assume a model for the stochastic process, which should guide the effort to estimate its characteristics.

The dissimilarity of the two transforms becomes particularly acute when considering observations as realizations of a stochastic process. For Fourier analysis, a stochastic process model exists: it is the generalized harmonic process (Priestley, 1981; Percival and Walden, 1993), a zero-mean, variance-stationary process (variance per unit of time is conserved). Without going into the details (e.g., Priestley, 1981; Percival and Walden, 1993; Papoulis and Pillai, 2002) we state that if the process *x*(*t*) is stochastically continuous and stationary, there exists a zero-mean process *χ*(*f*) with orthogonal increments such that

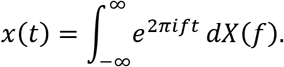

The existence of the spectral density of the process is guaranteed by the celebrated Wiener-Khinchin theorem (Priestley, 1981), which states that the power spectral density of process *x* is the Fourier transform of its auto-covariance function. It is important to note the condition of orthogonality of increments: it means that the Fourier components of different frequencies are uncorrelated. Therefore, the Fourier stochastic mode cannot have (does not include) non-stationary realizations such as sharp waves.

In contrast, as far as we are aware, there no widely-accepted stochastic-process model for transient signals. therefore, it appears that it makes little sense to look for a stochastic scope of the wavelet transform, as it is hard to imagine an overarching stochastic model for non-stationary processes (although recent attempts, e.g., Antoniou and Gustafson, 1999, might prove this argument wrong). While one is free to average whatever feature seems relevant, without a model for the stochastic process, we do not in fact know under what conditions a particular average exists.

It should be clear from the discussion above that the Fourier and wavelet representations are not interchangeable. For example, remapping a wavelet scalogram to frequencies and averaging over time to estimate the distribution of the power over frequencies has at least two fundamental problem: the frequency mapping is basically arbitrary, and the time average is not defined, if the process is non-stationary. Alternatively, if the process is stationary, the Fourier transform is the correct tool, because the wavelet distorts the distribution due to its peculiar shape of time-frequency atoms.

## A simple stationarity test

Remarkably, the simple idea that a variance-stationary generalized harmonic process should have at most weakly-correlated frequency components suggests a simple test for stationarity. The bicoherence of a stationary stochastic process should be weak (Elgar, 1987; Sheremet et al., 2016; Kovach et al., 2018). The examples shown in figure 13 illustrates this property. The stationary series, although weakly asymmetric (not entirely free of cross-spectrum correlation) exhibits a bicoherence that is essentially statistically zero, with the exception of the harmonic phase coupling. One could classify this series as matching the generalized harmonic stochastic process. In contrast, the non-stationary example in figure 13 shows large areas of strong coupling, suggesting that this should indeed be classified as non-stationary.

**Figure 13:**
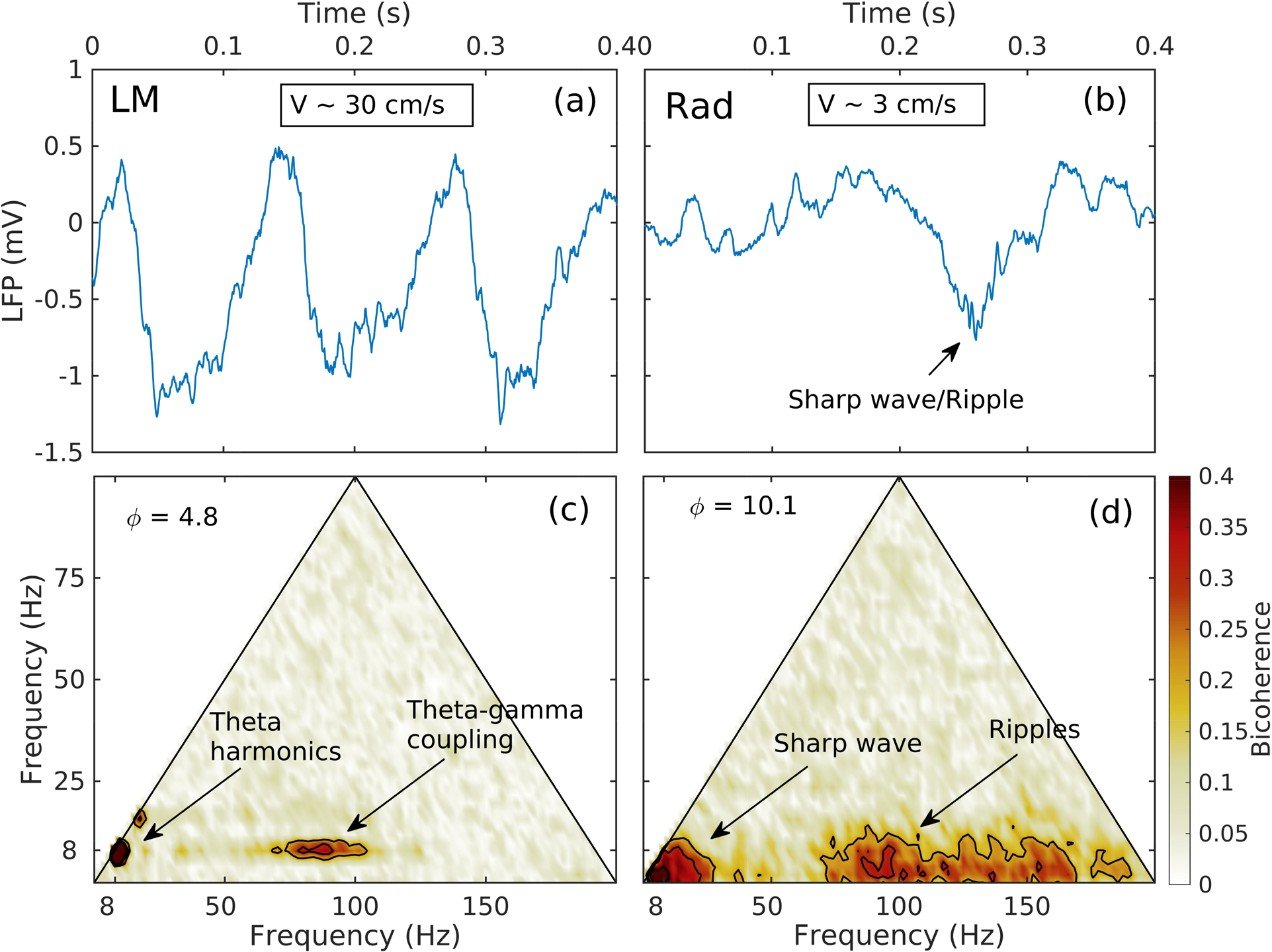
Examples of bicoherence estimates (c, d) for two LFP recordings (sample time series shown in panels a,b). Left: LM layer, active exploration at high speed motion; arrows mark the areas in the bicoherence map corresponding to theta/harmonics, and theta/gamma phase-coupling. Right: Rad layer, non-REM sleep; arrows mark the areas in the bicoherence map corresponding to the phase-coupling between Fourier components that assemble together into the solitary wave, and those that correspond to sharp wave and sharp-wave/ripples phase coupling.

## Acknowledgements

This work was supported by the McKnight Brain Research Foundation, and NIH grants-Grant Sponsor: National Institute on Aging; Grant number: AG055544 and Grant Sponsor: National Institute of Mental Health; Grant Number: MH109548 and a Diversity Supplement to NIH grant R01MH109548 (JPK). Special thanks to S.D. Lovett for technical support.

